# Mast cells are mediators of fibrosis and effector cell recruitment in dermal chronic graft-versus-host disease

**DOI:** 10.1101/668236

**Authors:** Ethan Strattan, Senthilnathan Palaniyandi, Reena Kumari, Jing Du, Natalya Hakim, Timothy Huang, Melissa V. Kesler, C. Darrell Jennings, Jamie L. Sturgill, Gerhard C. Hildebrandt

## Abstract

Allogeneic hematopoietic stem cell transplant (allo-HSCT) is often used to treat acute leukemia or defects of hematopoiesis. Its widespread use is hampered by graft-versus-host disease (GVHD), which has high morbidity and mortality in both acute and chronic subtypes. Chronic GVHD (cGVHD) occurs most frequently in skin and often is characterized by pathogenic fibrosis. Mast cells (MCs) are known to be involved in the pathogenesis of other fibrotic diseases. In a murine model of cGVHD after allo-HSCT, C57BL/6J recipients of allogeneic LP/J donor cells develop sclerodermatous dermal cGVHD which is significantly decreased in mast cell-deficient B6.Cg-Kit^W-sh/^HNihrJaeBsmGlliJ recipients. MCs survive conditioning and are associated with fibrosis, chemokine production, and recruitment of GVHD effector cells to the skin. Chemokine production by MCs is blocked by drugs used to treat cGVHD. The importance of MCs in skin cGVHD is mirrored by increased MCs in the skin of patients with dermal cGVHD. We show for the first time a role for MCs in skin cGVHD that may be targetable for preventive and therapeutic intervention in this disease.

## Introduction

Graft-versus-host disease (GVHD) has remained the major limiting factor for the success of allogeneic hematopoietic stem cell transplant (allo-HSCT) for decades. Chronic GVHD (cGVHD) is a subtype of GVHD which is a major contributor to patient morbidity and mortality, occurring in about 50 percent of patients after allo-HSCT^1^. It is a complex and enigmatic disease which can resemble autoimmune, inflammatory, or fibrotic immune responses, symptoms of which can occur in any organ system. Most frequently involved is the skin, with symptoms often presenting as sclerosis alongside increased collagen deposition^2^. Scleroderma-like cGVHD observed in transplant survivors can result in skin thickening and stiffness at localized sites or can manifest over extended areas of the body which can lead to full loss of mobility and entire body encasement.

Incomplete knowledge of the pathogenesis of cGVHD has long hampered therapeutic approaches. cGVHD is associated with risk factors related to acute GVHD and alloreactive donor immune cell responses. Perhaps because of this association, this debilitating disease is currently treated similarly to acute GVHD with the use of immunosuppressive and anti-inflammatory approaches, yet responses are not satisfactory. Furthermore, long-term treatment in this manner is associated with poor tolerability and risk of severe and potentially lethal infections^3^. Therefore, additional research into targetable effectors in cGVHD is a clinically unmet need.

Mast cells are long-lived, tissue-resident myeloid granulocytes, traditionally defined by their role in allergy through release of proinflammatory mediators such as histamine and heparin. This narrow role has been challenged in recent years, with studies pointing to novel roles for mast cells in wound healing, anti-venom, protection against bacterial pathogens, and recruitment of other immune cells^4^. Importantly, mast cells have been shown to be necessary for induction of pathogenic fibrosis in artificially-induced transgenic or bleomycin-induced models^5–7^, but this has not been studied in the context of cGVHD pathogenesis. Preliminary research on this topic was performed several decades ago although it has not since been thoroughly investigated^8–12^, likely because of the lack of effective models of mast cell deficiency and the need for better murine models of cGVHD. The development of mast cell-deficient mice and refinement of the murine models of allo-HSCT now allows for a more thorough investigation into the role of mast cells in cGVHD pathophysiology^13^.

In this study, we use a minor histocompatibility-mismatched murine model of allogeneic transplant to induce cGVHD of the skin by injection of LP/J bone marrow and splenocytes into either C57BL/6J (allo-WT) recipients or B6.Cg-Kit^W-sh^/HNihrJaeBsmGlliJ MC-deficient (allo-MCd) recipient animals. We find that WT mice develop symptoms of dermal cGVHD, while mast cell-deficiency results in markedly reduced symptomology. Further, we show that mast cells are radioresistant and that recipient-derived mast cells are present in allo-WT post-transplant. These mast cells correlate with increased chemokine production and immune effector cell infiltration. Mast cell derived chemokine production was inhibited by ibrutinib or ruxolitinib, both of which are used to treat dermal cGVHD in patients.

This study is the first of its kind to pinpoint mast cells *in vivo* as effectors of fibrosis and effector cell recruitment in dermal cGVHD. Targeting of these tissue-specific cells could allow for reduction of cGVHD symptoms, and we hypothesize that the efficacy of current treatments for cGVHD could be due in part to blockade of mast cell responses in cGVHD target organs.

## Results

### Bone marrow-derived mast cells survive and are functional after ionizing radiation

Previous literature by several groups indicated that mast cells might be capable of surviving high doses of ionizing radiation, such as that used in conditioning prior to transplant of donor cells^8,9,14^. To determine this, we derived mast cells *ex vivo* from murine bone marrow precursors from both BALB/cJ and C57BL/6J mice. After 4-6 weeks of culture, these cells displayed morphological similarity to mast cells, and at least 97 percent displayed the characteristic markers of mast cells, FCeR1a/cKit +/+ by flow cytometry (Fig. 1A).

**Figure 1:**
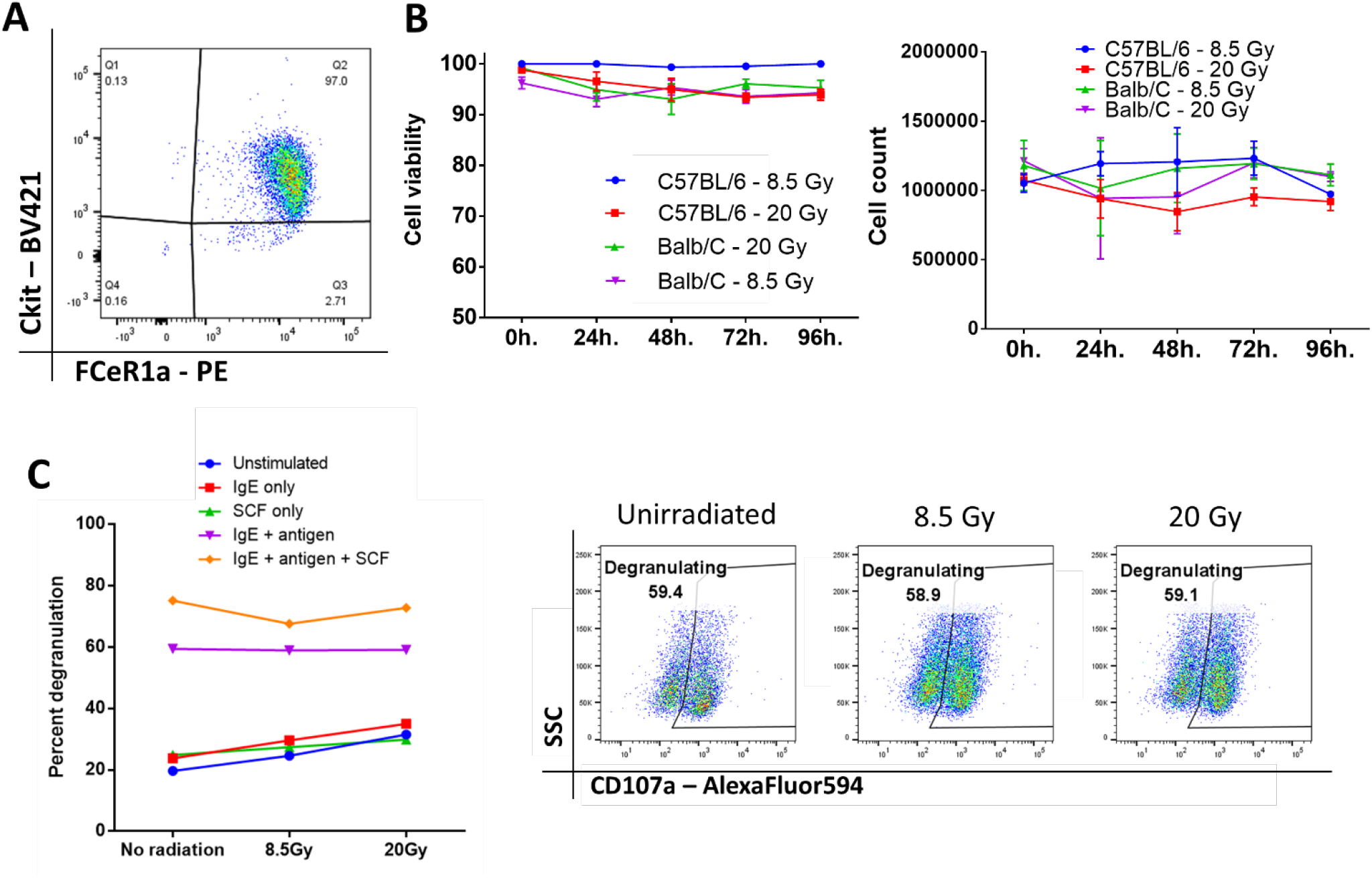
Bone marrow-derived mast cells survive and are functional after ionizing radiation. A) Phenotypic analysis of bone marrow-derived mast cells after 4-6 weeks of culture, displaying marker expression characteristic of mast cells and high purity. B) Mast cell viability and count after ionizing radiation. There was no significant change from baseline even 96 hr. after irradiation. Graphs are combined data from 3 independent experiments. C) Mast cells are still functional after irradiation. There is no significant difference between any of the doses across any group. Graph and flow plots are representative of two independent experiments. P value 0.01 to 0.05 = *, P value 0.001 to 0.01 = **, P value 0.0001 to 0.001 = ***, P value <0.0001 = ****, NS = not significant

We then tested the viability and overall number of these cells after they were treated with either 8.5 or 20 Gy of ionizing radiation. Mast cells from either C57BL/6J or BALB/cJ mice showed no change from baseline after 96 hours of culture, implying that mast cells are highly resistant to cell death induced by ionizing radiation (Fig. 1B). We then tested whether these cells would remain functional after irradiation. Degranulation in response to FCeR1a receptor crosslinking was measured by CD107a expression on the cell surface. Degranulation was apparent at all doses and was unaffected by ionizing radiation (Fig. 1C).

### Mast cell-deficient mice have improved outcome and decreased skin fibrosis after transplant

Having seen that mast cells are viable and functional after irradiation, we induced cGVHD *in vivo* through a minor histocompatibility mismatched murine bone marrow transplant model (LP/J → C57BL/6J)^15^, illustrated in Figure 2A. Allogeneic bone marrow transplants were performed into both wild type C57BL/6J (allo-WT) and B6.Cg-Kit^W-sh^/HNihrJaeBsmGlliJ mast cell-deficient mice (allo-MCd). Syngeneic transplant recipients did not develop cGVHD and served as a negative controls. At 7 weeks post-transplant, 18.75% of the allo-WT group had died from cGVHD, while there were no deaths in either the syngeneic or the allo-MCd mice (Fig. 2B). Using a previously-published whole-animal GVHD scoring system^16^, we saw significant decreases in GVHD scoring in allo-MCd mice as compared to allo-WT mice by week 5 (Fig. 2C).

**Figure 2:**
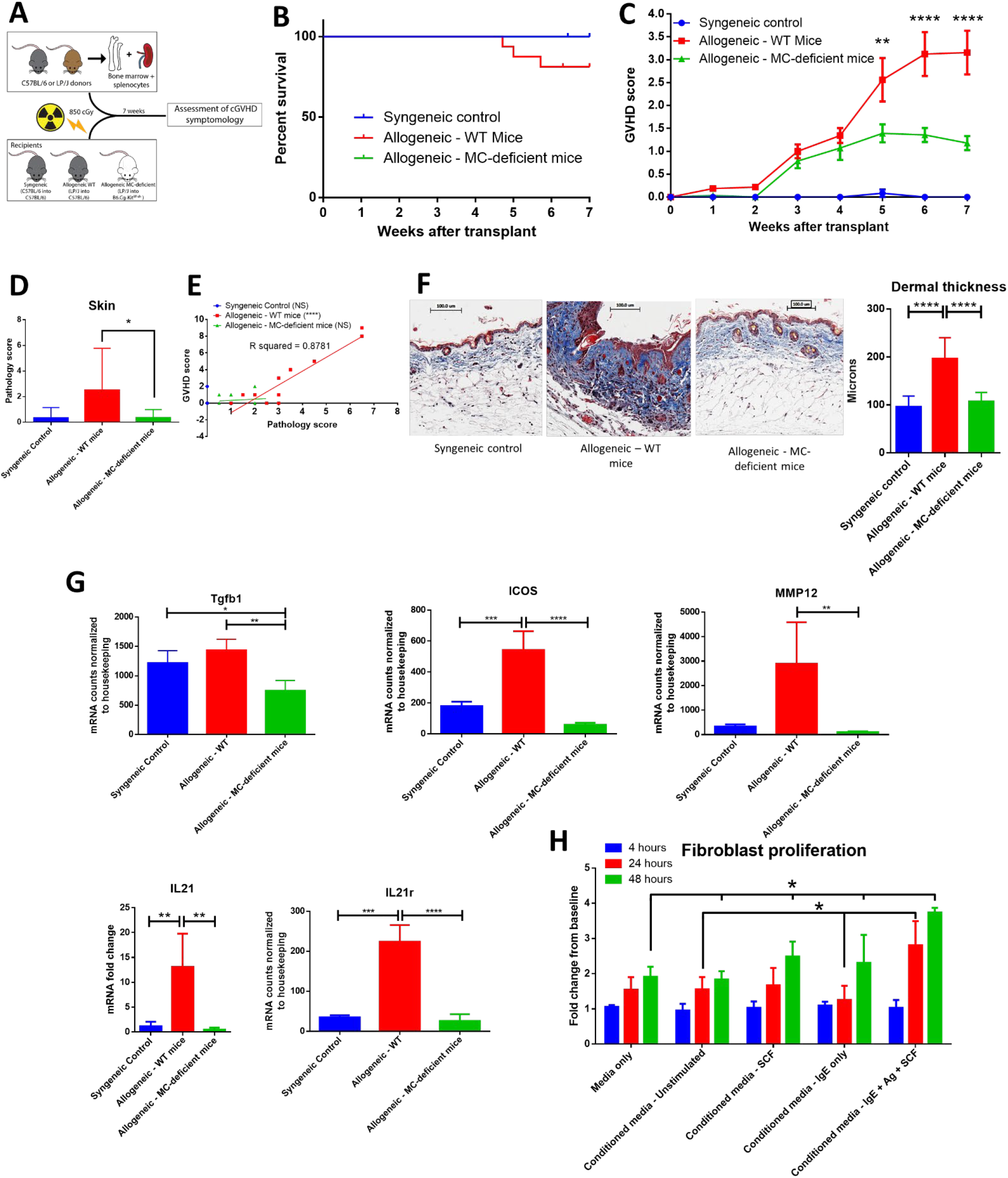
Mast cell-deficient mice have improved outcome, including less skin pathology and decreased fibrosis after transplant. A) Schematic of transplant to test the effects of mast cell-deficiency in cGVHD. B) Kaplan-Meier survival curve of syngeneic (n=6), allo-WT (n=16), and allo-MCd (n=14) groups after transplant. The curves are not significantly different, though the syngeneic and allo-MCd groups had no deaths, while the allo-WT group had 3 deaths. Data is combined from two independent transplants. C) Weekly GVHD scoring of syngeneic (n=6), allo-WT (n=16), and allo-MCd (n=14) groups after transplant. By week 5, there is a significant reduction in GVHD score in allo-MCd mice as compared to allo-WT mice as measured by a repeated-measures one-way ANOVA. Error bars are the standard error of the mean (SEM). D) Skin pathology score is significantly decreased in the allo-MCd (n=16) group as compared to the allo-WT group (n=14). Scoring accounts for damage to the epidermis, dermis, subcutaneous fat, and hair follicles, as well as inflammatory infiltrate and was performed by a blinded pathologist. Data is combined from two independent transplants. E) Whole-animal GVHD scoring significantly correlates with skin pathology scoring. R squared and P values were determined by Spearman nonparametric correlation. F) Representative images and quantification of Masson’s trichrome staining of skin from syngeneic (n=6), allo-WT (n=16), and allo-MCd (n=14) groups after transplant. Mice were quantified for dermal thickness according to the height of the blue collagen band across 10-20 different fields per mouse. Images are mice with dermal thickness closest to the mean values for each group. Quantification is combined data from two independent transplants. G) Genes associated with dermal fibrosis are significantly upregulated in allo-WT animals, as measured by NanoString analysis or qPCR (IL21 only). Samples were skin from syngeneic (n=3), allo-WT (n=3), and allo-MCd (n=5) mice. H) Supernatants from activated mast cells cause increased proliferation in fibroblasts. 48 hours after plating, supernatants from mast cells stimulated with IgE + antigen + 50ng/ml SCF caused fibroblasts to proliferate significantly more than any other condition. Data is representative of two independent experiments. P value 0.01 to 0.05 = *, P value 0.001 to 0.01 = **, P value 0.0001 to 0.001 = ***, P value <0.0001 = ****, NS = not significant

The mice were sacrificed and analyzed after week 7 post-transplant. Syngeneic mice had almost no skin pathology. However, allogeneic transplantation induced severe skin pathology in WT mice, which was significantly ameliorated in allo-MCd mice, as scored by an independent, blinded, board-certified pathologist (hematoxylin and eosin [H+E] staining, Fig. 2D). In the lung, small intestine, liver, and colon, there were no significant pathologic differences between the allogeneic groups (Supplementary Fig. 1). In the allo-WT but not the syngeneic or allo-MCd animals, pathology scores significantly correlated with clinical GVHD scores (Fig. 2E). Only mice in the allo-WT group displayed clinical symptoms consistent with sclerotic chronic GVHD, including scaly skin, hair loss, and patchy lesioning^17^. When skin from these mice was examined histologically by Masson’s trichrome staining, fibrotic cGVHD was evident only in the allo-WT group, including excess collagen deposition, hyperkeratosis, epidermal disruption, and significant increases in dermal thickness^18^, as shown by representative images and the corresponding quantification in Figure 2F. Importantly, no cGVHD was noted in the allo-MCd mice.

We next analyzed RNA extracted from skin of mice from each group by the NanoString Myeloid Innate Immunity panel or by qPCR for IL21 (not included in the NanoString panel). Significant increases in genes associated with skin fibrosis were evident in the allo-WT group, each of which was significantly lowered in the allo-MCd group (Fig. 2G)^19,20^. Taken alongside the histologic data, this indicates a mast cell-dependent induction of fibrosis through an unknown mechanism.

Since fibroblasts are the main effector cell responsible for collagen deposition and fibrosis, we assessed whether supernatants from mast cells could affect the proliferation of fibroblasts^21^. We isolated and cultured fibroblasts derived from the skin of C57BL/6J mice for 2-3 weeks. Following trypsinization, these cells were transferred to a 6 well plate. After 4 hours to allow cells to adhere, there were no differences in count between groups. We then added fresh media or conditioned media from mast cells stimulated under various conditions to these cultures and counted them after 24 and 48 hours. We saw that after 24 hours, fibroblasts cultured in conditioned media from activated mast cells (IgE + antigen + SCF) had grown significantly more than fibroblasts incubated with conditioned media from unstimulated mast cells or from mast cells incubated with IgE but without antigen. By 48 hours post-plating, fibroblasts cultured in conditioned media from activated mast cells showed a significant increase in cell count compared to all other conditions (Fig. 2H). This indicates that products from activated mast cells can induce fibroblast proliferation which could lead to increased fibrosis *in vivo*.

### Mast cells survive conditioning and are recipient-derived after murine bone marrow transplant

During a bone marrow transplant, the donor stem cells reconstitute and fill the role previously played by the recipient bone marrow. Among these roles are hematopoiesis of lymphoid and myeloid immune subsets, mast cells included^1^. *Ex vivo*, we have shown that mast cells can survive radiation, but we wanted to determine if mast cells survived post-transplant *in vivo;* in addition to radiation, recipient-derived mast cells could also be eliminated by alloreactive donor T-cells. Therefore, we examined tissue sites where mast cells are tropic, including skin, ear, and trachea, as well as other GVHD target organs (small intestine, lung and liver).

Seven weeks after transplant, organ tissue was stained with toluidine blue and mast cells were counted per high-power field (400x). Skin, ear, and tracheal tissue showed a similar pattern, with low numbers of mast cells in syngeneic animals, higher numbers in allo-WT animals, and low-to-undetectable levels of mast cells in allo-MCd animals (Fig 3A, 3B, 3C). Mast cells were low-to-undetectable in small intestine, lung, and liver and there were no differences between groups (data not shown). Findings were verified by a blinded, independent researcher. To confirm our finding in the skin, we returned to our NanoString data set, where we found that mast cell-specific genes including chymase, tryptase, FCeR1a, and Cpa3 were expressed at extremely low levels in the skin of allo-MCd mice (Fig. 3D).

**Figure 3:**
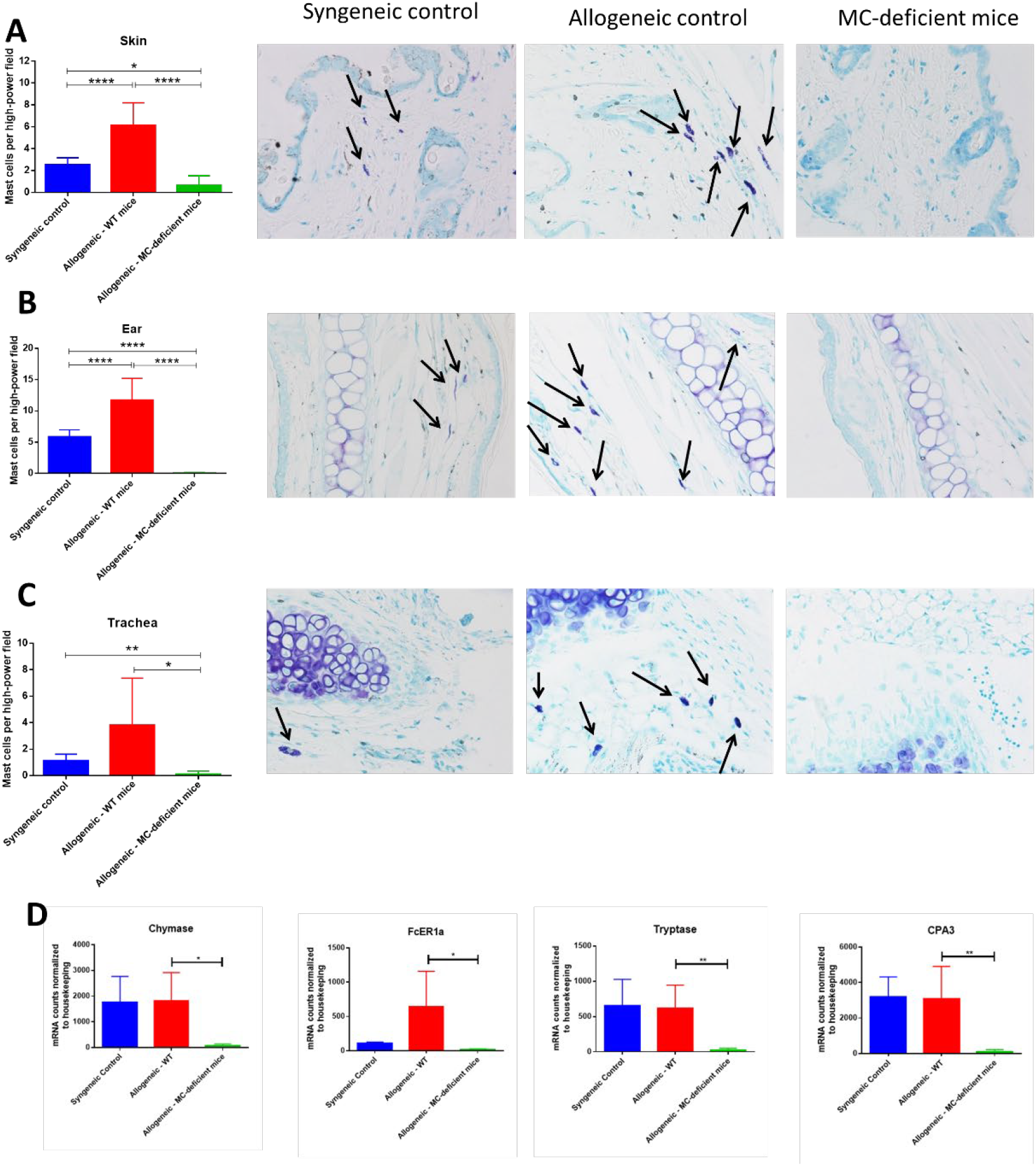
Mast cells survive conditioning and are recipient-derived after murine bone marrow transplant. Mast cell counts and toluidine blue-stained representative images in the A) Skin, B) Ear, and C) Trachea of syngeneic (n=6), allo-WT (n=16), and allo-MCd (n=14) animals. Mast cell counts per high-power field (400x) are displayed in the first column alongside images from animals whose mast cell count was nearest the mean value for the group. Mast cells were denoted by metachromatic staining and granular appearance. The lack of mast cells in the allo-MCd group implies that mast cells present in the other groups survived transplant and are recipient-derived, rather than having been reconstituted from donor-derived bone marrow precursors. D) Levels of mast cell-specific genes in the skin are significantly lowered in allo-MCd mice. Data displayed is from syngeneic (n=3), allo-WT (n=3), and allo-MCd (n=5) mice, as measured by NanoString analysis. P value 0.01 to 0.05 = *, P value 0.001 to 0.01 = **, P value 0.0001 to 0.001 = ***, P value <0.0001 = ****, NS = not significant

The extremely low numbers of mast cells in the allo-MCd compared to our allo-WT group, even 7 weeks post-transplant, implies that mast cells were unable to fully reconstitute from the bone marrow into their tissue niche. Importantly, MC-deficient mice are not defective in their capacity to support mast cell growth as shown in re-engraftment experiments^22^. The mast cells present in syngeneic and allo-WT animals therefore are likely mainly recipient-derived and had survived conditioning and the post-transplant environment.

### The dermal environment of allo-WT mice is enriched in chemokine signaling compared to allo-MCd mice

Our findings demonstrate a difference in GVHD scoring and skin pathology, yet given the pleotropic effects of mast cells in the immune system we next decided to explore their functional role. We examined immune cells in the spleen and saw no differences in proportion or count in myeloid or lymphoid subsets (Supplementary Fig. 2). To examine more local factors, skin from animals from each group was analyzed using the NanoString Myeloid Innate Immunity panel. When we looked at the subset of genes overexpressed in allo-WT vs. allo-MCd by PANTHER pathway analysis^23^, we found only one pathway to be statistically significant, an approximately 5-fold enrichment for the “Inflammation mediated by chemokine and cytokine signaling pathway” (PANTHER pathway number P00031, Supplementary Fig. 3A). Cytokine levels were found to be expressed at low levels in both skin and serum and were largely unchanged between groups (Supplementary Figure 3B-D).

Further analysis was performed using the NanoString nSolver software which clusters genes by pre-annotated pathways. Sorting on “Chemokine signaling genes,” heatmap analysis demonstrated an overall trend towards increased dermal expression of genes involved in these pathways in allo-WT mice, with lower expression in allo-MCd mice (Fig. 4A). We further examined expression of specific chemokine transcripts. CCL2, CCL3/CCL4 (which encode MIP-1a and MIP-1b, respectively), and CCL5 all demonstrated significant upregulation of transcript in the skin in allo-WT mice (Fig. 4B). When we measured protein levels of these chemokines by cytometric bead array, we saw an extremely similar expression pattern, with all 4 chemokines upregulated in allo-WT, while syngeneic and allo-MCd were not significantly different (Fig. 4C). Intriguingly, the levels of these chemokines in the allo-WT group were all significantly correlated with their overall GVHD score (Fig. 4D). All of these chemokines can be chemotactic for T-cells, the canonical GVHD effector cell^24,25^. Additionally, they are capable of recruiting monocytes, macrophages, or other myeloid cells, many of which are implicated in cGVHD pathogenesis and fibrotic disease.

**Figure 4:**
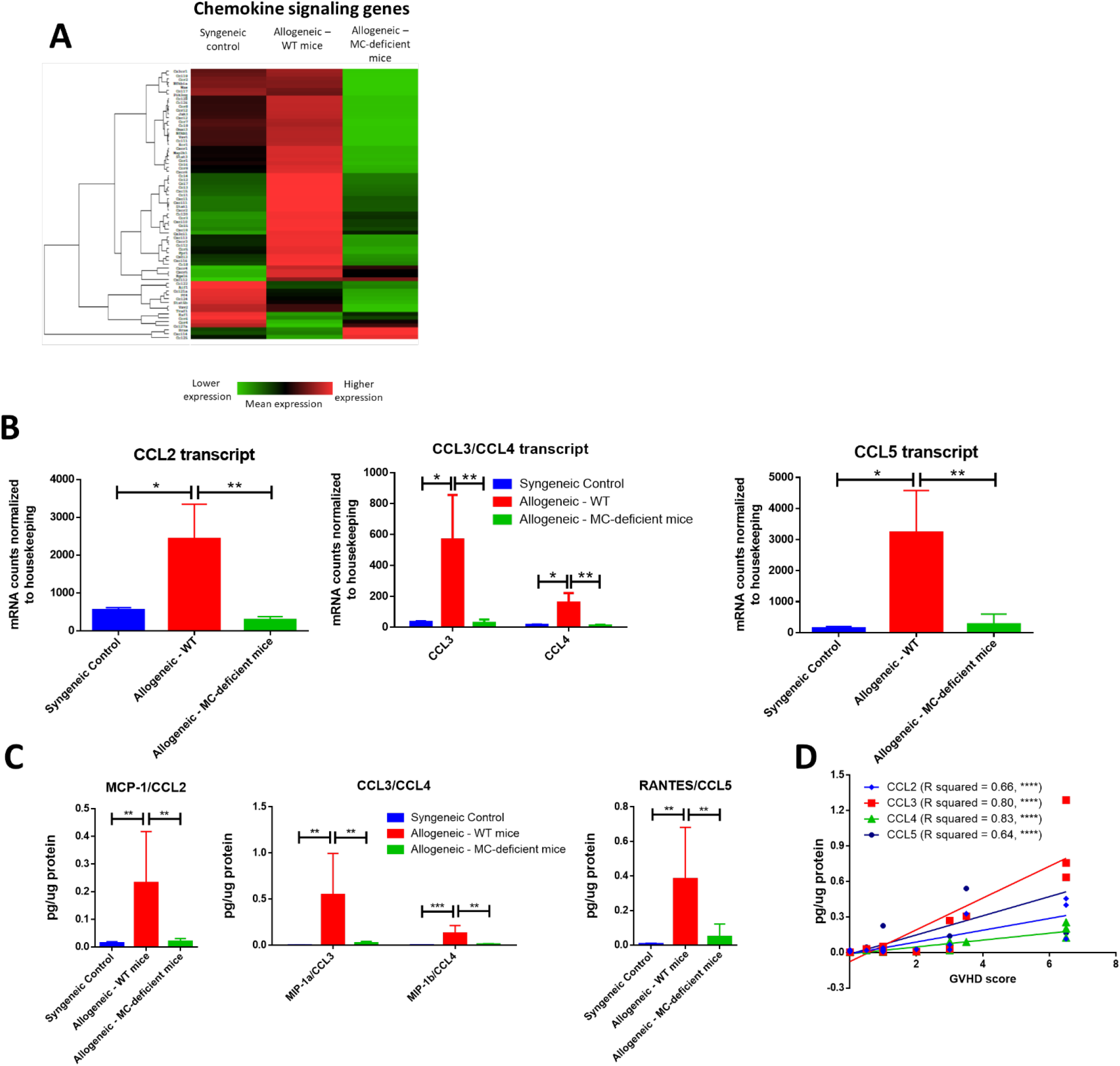
The dermal environment of allo-WT mice is enriched in chemokine signaling compared to allo-MCd mice. A) Heatmap analysis showing lowered expression of chemokine signaling genes in allo-MCd animals compared to allo-WT animals as measured by NanoString. Heatmaps and gene pathway annotations were generated using NanoString nSolver software. B) Transcript as measured by NanoString, and C) protein levels of chemokines are upregulated in the skin of allo-WT mice, while syngeneic mice and allo-MCd mice are not significantly different. (syngeneic n=6, allo-WT n=6, and allo-MCd n=8, data are combined from two independent transplants). Protein levels were measured by LEGENDplex Chemokine Cytometric Bead Array kit. All NanoString data displayed is from syngeneic (n=3), allo-WT (n=3), and allo-MCd (n=5) mice. D) Protein levels of CCL2, CCL3, CCL4, and CCL5 all significantly correlate with whole-animal GVHD scoring. R squared and P values were determined by Spearman nonparametric correlation. P value 0.01 to 0.05 = *, P value 0.001 to 0.01 = **, P value 0.0001 to 0.001 = ***, P value <0.0001 = ****, NS = not significant

### Immune infiltration and activation are ameliorated by mast cell deficiency

Given the inflammation and increased chemokines evident in the skin of our allo-WT mice, we wanted to determine whether this resulted in increased immune infiltration and tissue damage. Indeed, when we examined H+E-stained skin sections from allo-WT mice, epidermal dysregulation and an obvious dermal immune infiltrate was visible (Fig. 5A, quantified in Fig. 5B). Animals from the syngeneic and allo-MCd groups displayed equally low levels of immune infiltrate. The immune infiltrate was characterized as to its hematopoietic lineage and was found to be predominantly lymphocytic in all three groups (Fig. 5C). Accordingly, there was no significant difference across all groups in eosinophil or neutrophil count in H+E-stained skin sections, nor was there a difference in CD19 transcript by qPCR (Supplementary Fig. 2C). All characterization and counting on H+E slides were performed by an independent, board-certified pathologist.

**Figure 5:**
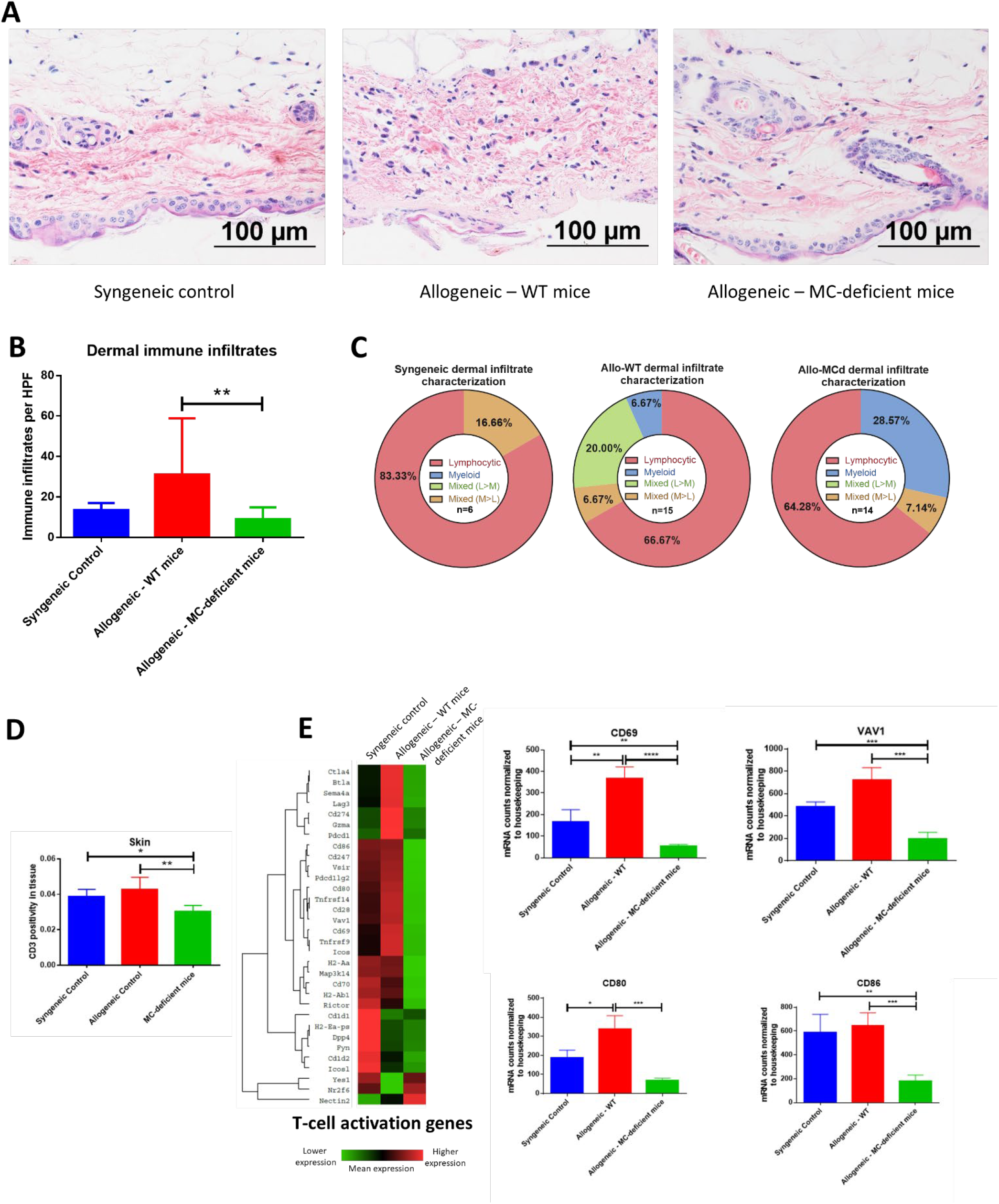
Immune infiltration and activation are ameliorated by mast cell deficiency. A) Representative images from H+E staining show an increased immune infiltrate and tissue damage in skin from allo-WT animals, while syngeneic and allo-MCd animals have a similar phenotype. In allo-WT animals, epidermal dysregulation and an obvious dermal immune infiltrate is visible. B) Quantification of Figure 5A. C) The immune infiltrate was characterized as to its hematopoietic lineage by morphology and was found to be predominantly lymphocytic in all three groups. D) Immunohistochemistry against CD3 demonstrates a significant increase in T-cell infiltration in the skin of allo-WT as compared to allo-MCd animals. Slides were digitally scanned using the Aperio ScanScope XT, then quantified for DAB chromogen staining across the whole tissue using the Positive Pixel Count V9 algorithm. E) Heatmap and gene-level analysis demonstrates significant increases in genes involved in the T-cell activation pathway in allo-WT as compared to allo-MCd animals as measured by NanoString. This indicates that despite a slight but significant increase in T-cells, activation status of those cells widely differs between groups. Heatmaps and gene pathway annotations were generated using NanoString nSolver software. NanoString data displayed is from syngeneic (n=3), allo-WT (n=3), and allo-MCd (n=5) mice. P value 0.01 to 0.05 = *, P value 0.001 to 0.01 = **, P value 0.0001 to 0.001 = ***, P value <0.0001 = ****, NS = not significant

T-cells in the skin, as measured by immunohistochemistry against CD3, were upregulated from allo-MCd to allo-WT animals (Fig. 5D). This slight but significant increase in cellularity was seen alongside increased gene expression of markers of T-cell activation including CD69 and VAV1, as well as markers of costimulation such as CD80 and CD86 (Fig. 5E). It seems that while the level of T-cell infiltration is not hugely different, activation and the costimulatory milieu in the dermal environment may enhance their effects.

### Mast cells produce chemokines upon stimulation which are blocked by drugs used in treatment of cGVHD

Mast cells are known to be able to produce a wide variety of mediators, as reviewed excellently by Mukai *et al* ^26^. As shown in Figure 4D, chemokine production seems to be a major factor in the pathogenesis of sclerodermatous GVHD in this model. Therefore, we wanted to test whether mast cells were capable of producing these same cytokines, assayed *ex vivo* using bone marrow-derived mast cells cultured from C57BL/6J mice.

Indeed, we saw that upon stimulation with multiple methods of mast cell activation (IgE/antigen crosslinking or IgE/antigen + IL-33), mast cells produced significant amounts of CCL2, CCL3, and CCL4. Importantly, treatment with ibrutinib, a BTK inhibitor, or ruxolitinib, a Jak1/Jak2 inhibitor, reduced mast cell production of these chemokines in a dose-dependent manner (Fig. 6A, 6B, 6C). Treatment with imatinib or fingolimod did not reduce mast cell chemokine production even at the highest doses tested (Supplementary Fig. 4A). Drug dosing had no effect on cell viability (Supplementary Fig. 4B). While CCL5 was the other chemokine significantly upregulated in the skin of allo-WT mice (Fig. 4D), we saw no CCL5 production from mast cells in this context, though previous papers from our group have pointed out donor-derived T-cells as the primary producers of CCL5 in the post-transplant setting^27^.

**Figure 6:**
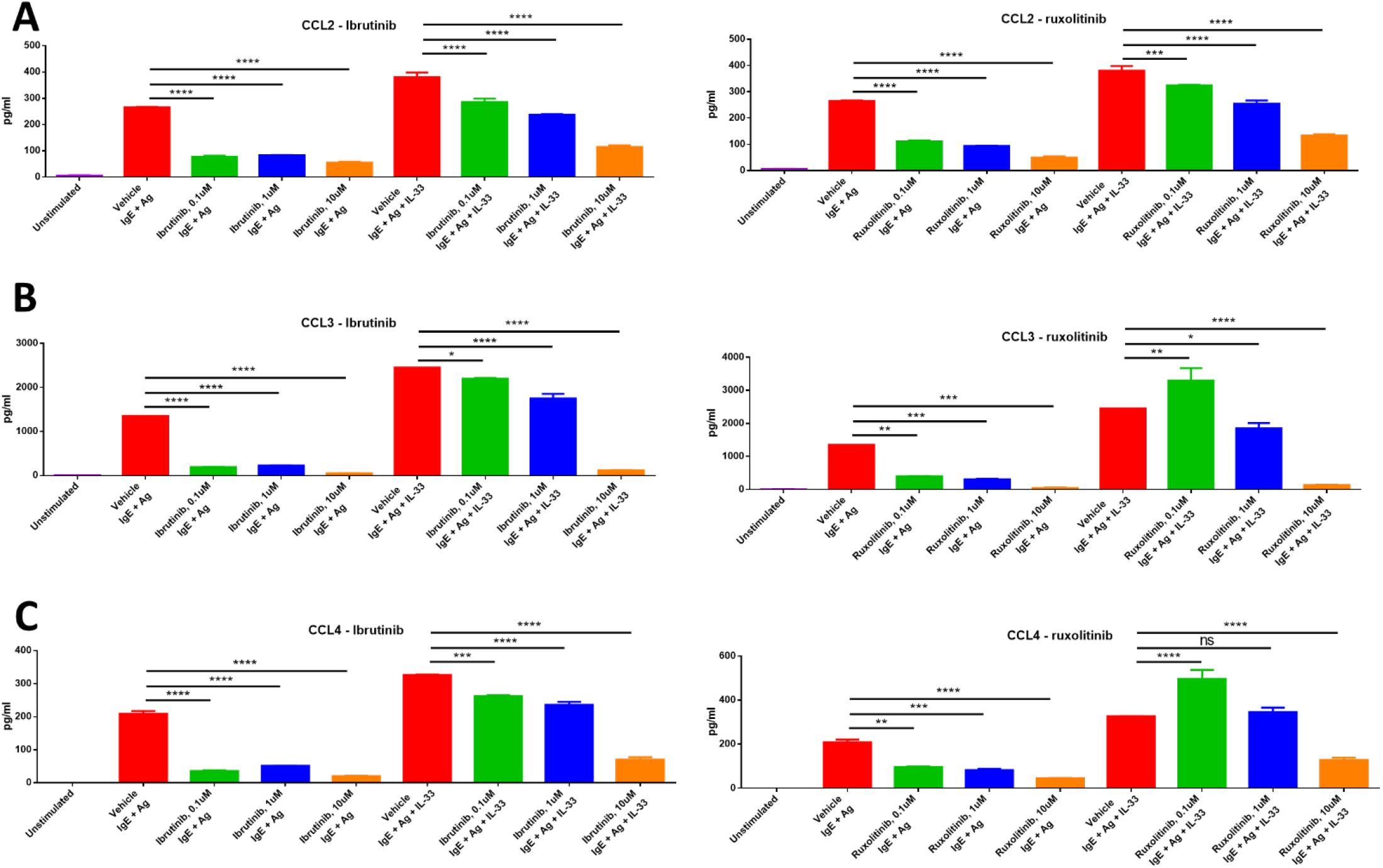
Mast cells produce chemokines upon stimulation, production of which is blocked by drugs used in treatment of cGVHD. Mast cells produce high levels of A) CCL2, B) CCL3, and C) CCL4 upon stimulation with IgE + antigen or IgE + antigen + IL-33 (column 1 vs. columns 2 and 6). Production of these chemokines is inhibited by treatment with either ibrutinib or ruxolitinib in a dose-dependent manner. Results shown are representative of 2-4 independent assays. Error bars are the SD of technical replicates. Chemokine assays were performed using the LEGENDplex Inflammatory Chemokine Assay kit, which measures levels of 13 chemokines. Mast cells did not produce significant amounts of CCL5, CCL11, CCL17, CXCL1, CXCL9, CXCL10, CXCL13, CXCL5, or CCL22 (data not shown). P value 0.01 to 0.05 = *, P value 0.001 to 0.01 = **, P value 0.0001 to 0.001 = ***, P value <0.0001 = ****, NS = not significant

### Mast cell numbers are increased in the skin of patients with dermal manifestations of cGVHD

To further examine this phenomenon in the context of human patients with cGVHD, we obtained approval from the University of Kentucky Institutional Review Board for studies on banked, deidentified human samples (see methods for details). We used the University of Kentucky Biospecimen Procurement and Translational Pathology Shared Resource Facility (BPTP SRF) to identify skin biopsies from allo-HSCT recipients who were diagnosed with chronic GVHD and where histologic changes were consistent with dermal graft-versus-host disease. Samples from 5 cGVHD patients were stained with Masson’s trichrome alongside healthy patient skin. Epidermal thickness was measured by Masson’s trichrome staining and found to be increased due to collagen deposition, consistent with cGVHD symptomology (Fig. 7A). We identified levels of tryptase-positive cells in the skin by immunohistochemistry in order to quantify the number of mast cells present in these samples. Tryptase-positive cells were primarily located near the dermal-epidermal junction and were significantly increased in cGVHD patients as compared to normal skin samples as measured by Aperio (Fig. 7B). Quantification was verified by a blinded independent researcher. Representative images for the staining are shown in Fig. 7C. These data correlate with our findings that mast cells may be important in the pathogenesis of dermal chronic graft-versus-host disease.

**Figure 7:**
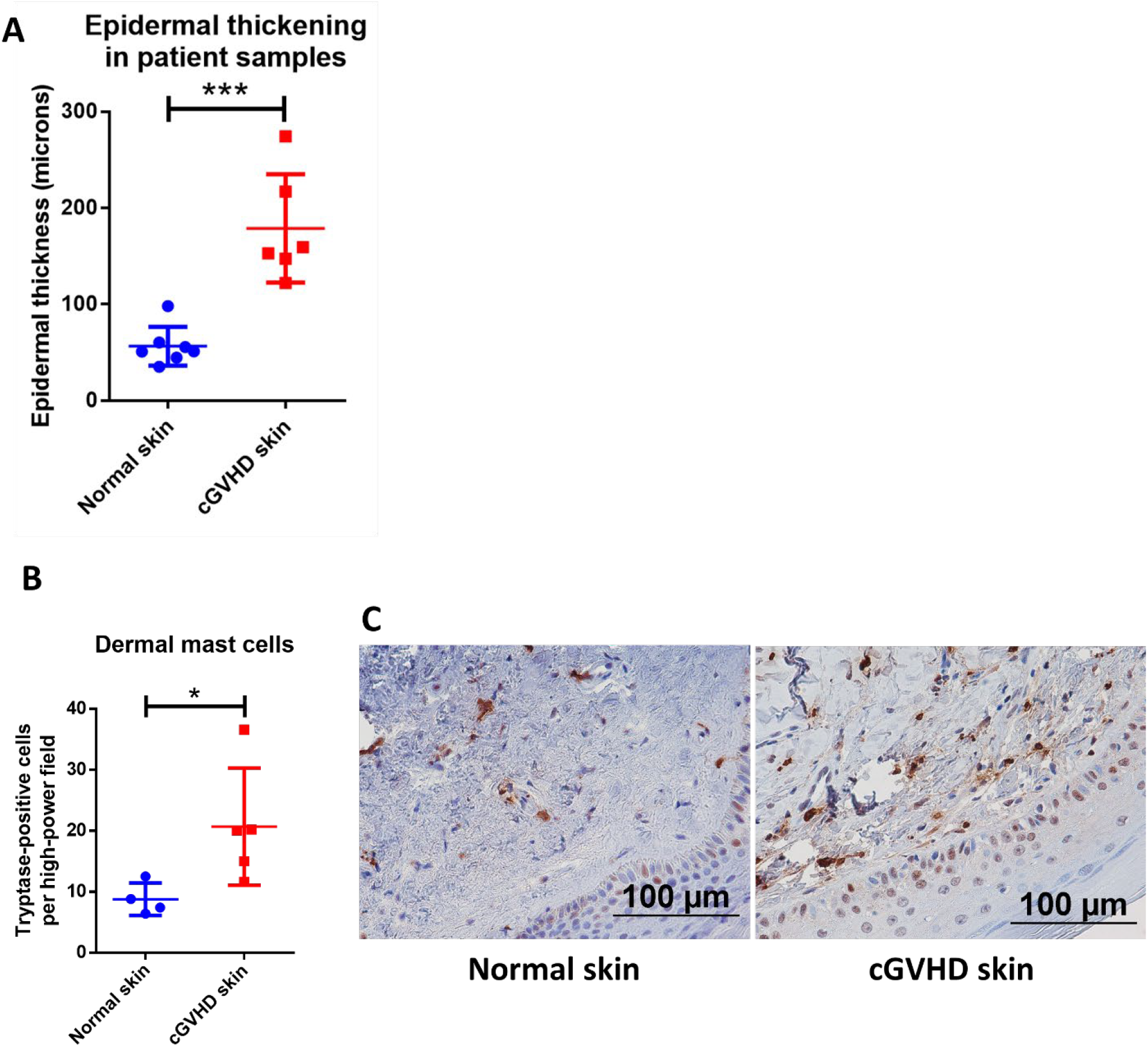
Mast cell numbers are increased in the skin of patients with dermal manifestations of cGVHD. A) Skin biopsies from patients with cGVHD displayed histologic changes consistent with dermal cGVHD including epidermal thickening as compared to normal controls (normal controls n=4, cGVHD patients n=5). B) cGVHD patient samples had a significant increase in tryptase staining as measured by Aperio ImageScope in the dermis as compared to normal controls (normal controls n=4, cGVHD patients n=5). P value 0.01 to 0.05 = *, P value 0.001 to 0.01 = **, P value 0.0001 to 0.001 = ***, P value <0.0001 = ****, NS = not significant

## Discussion

Pathogenic fibrosis is a characteristic of many diseases in the developed world, with up to 45 percent of all deaths associated with symptoms of chronic fibrosis^21^. Understanding the mechanisms behind fibrosis and developing new treatments is a major public health concern. As in other fibrotic diseases, treatment of cGVHD is hampered by incomplete knowledge of the mechanisms driving disease pathogenesis. Many of the current treatments, such as immunosuppressive steroids, do not bring about the desired patient outcomes and are unsuited for the long-term course of treatment often required in cGVHD. In this study, we demonstrate a novel role for mast cells in the pathogenesis of dermal cGVHD and show the first evidence of mast cell involvement *in vivo*, which could open new paradigms in the treatment of fibrotic cGVHD.

Our data demonstrating mast cell viability and functionality after radiation confirms several earlier studies showing similar results(Fig. 1B, 1C)^14,28^. This radioresistance of mast cells is striking in contrast to many other immune cell types, which are considered to be depleted during the conditioning phase of allo-HSCT^17^. Taken together, this implies that the mast cell is a radioresistant, mature, immune effector cell, which remains functional and that has tropism towards the cGVHD target organs. It is therefore perhaps unsurprising that there was such a dichotomous difference in cGVHD symptomology *in vivo* between the allo-WT and allo-MCd groups in our murine model of allogeneic transplant. Using the LPJ→C57BL/6J model of murine cGVHD in both WT C57BL/6J and B6.Cg-Kit^W-sh^/HNihrJaeBsmGlliJ MC-deficient animals, we showed evidence for reductions in whole-animal GVHD scoring, skin pathology, and in dermal thickness and fibrosis in the allo-MCd mice. This correlated with decreases in transcript level of genes associated with fibrosis in the dermal environment (Fig. 2A-G)^19,20^. Additionally, our *in vitro* evidence demonstrated that mast cells are capable of producing factors that induce proliferation in skin-derived fibroblasts, which are responsible for collagen deposition (Fig. 2H). Mast cells are known to be one of the major sources of fibroblast growth factor, which could explain this phenomenon^29^.

Previous work by Leveson-Gower *et al.* has shown that the presence of mast cells is beneficial in the context of acute GVHD through mast cell production of anti-inflammatory factors, such as IL-10^30^. However, that study was performed in a C57BL/6→BALB/c model of acute GVHD, where defining characteristics of the disease include acute inflammation and early alloreactive damage to the tissues, along with rapid disease onset, progression and mortality from acute GVHD. In contrast, chronic GVHD is often defined by inappropriate and prolonged inflammation alongside anti-inflammatory “wound healing” responses and fibrosis^31^. Therefore, we believe that the mast cell, while beneficial in the inflammatory phase of aGVHD, ultimately becomes detrimental in cGVHD through its ongoing anti-inflammatory activities resulting in collagen deposition and pathogenic fibrosis.

These data fit with a study by Levi-Schaffer *et al.*, which demonstrated that mast cells are activated by splenic supernatants from mice with cGVHD, as well as their work showing that fibroblasts from mice with cGVHD enhance proliferation of connective tissue-derived mast cells^10,12^. Taken together with our data showing that mast cell products can stimulate proliferation of fibroblasts (Fig. 2H) this could indicate a positive feedback loop in cGVHD wherein mast cells and fibroblasts are able to induce proliferation in the other cell type. In addition, there is a growing body of literature in human and murine models demonstrating that mast cells and fibroblasts have both direct and indirect interactions, the result of which is increased collagen deposition and fibrosis^5,32–34^.

Our data also demonstrated that *in vivo*, recipient-derived mast cells survive at least 7 weeks post-transplant. Mast cell-deficiency was maintained in the allo-MCd group, while the syngeneic and allo-WT groups still had significant numbers of mast cells, as measured by toluidine blue staining. This finding was confirmed by transcript-level analysis of mast cell-specific genes (Fig. 3). We further saw that the allo-WT animals had significant increases in transcript and protein levels of chemokines (CCL2, CCL3, CCL4, and CCL5) that can recruit canonical cGVHD effector cells. Levels of these chemokines significantly correlated with our whole-animal GVHD scoring (Fig. 4). Additionally, the skin of allo-WT mice had increased tissue damage and a primarily lymphocytic immune infiltrate. We also showed a concomitant increase in T-cell number by immunohistochemistry, alongside data demonstrating increased levels of genes involved in T-cell activation and signaling (Fig. 5).

Importantly, all these changes were ameliorated by mast cell-deficiency. Given the mast cell’s well-known role as a “sentry cell^35^,” it seems that in the context of cGVHD pathogenesis they may be necessary for initial recruitment of inflammatory cells which then cause a feedback loop of further inflammation and immune recruitment. Additionally, they may have more direct effects through their impact on fibroblast proliferation. We also show that mast cells are capable of production of many of the chemokines that are upregulated in the allo-WT animals (Fig. 6A, B, C, **columns 1, 2, 6**).

Ibrutinib is a combined BTK/ITK inhibitor known to be efficacious in treatment of murine and human cGVHD, for which it is currently the only FDA-approved drug^36,37^. The Jak1/Jak2 inhibitor, ruxolitinib has been in case studies and is currently in Phase 2 and 3 clinical trials for use in steroid-refractory cGVHD^38–40^. Mast cells express many of the signaling molecules targeted by these drugs, and we therefore investigated whether chemokine production would be inhibited when treated with ibrutinib or ruxolitinib^41,42^. Indeed, we found a dose-dependent decrease in production of CCL2, CCL3, and CCL4 when treated with ibrutinib and ruxolitinib (Fig. 6). This phenomenon was not seen in cells treated with imatinib or fingolimod, which is unsurprising given the lack of exogenous S1P or SCF in the assay (Supp. Fig. 4). Given the overexpression of these same chemokines in the skin of the allo-WT vs. allo-MCd mice (Fig. 4E) and the reduction in skin symptomology of patients treated with ibrutinib and ruxolitinib^38^, it may be that inhibition of mast cell chemokine production is a component of the mechanism by which these inhibitors are clinically efficacious.

Lastly, we show that patients with dermal manifestations of cGVHD have higher levels of mast cells in the skin than normal controls (Fig. 7). This data is similar to that seen in our murine model and is suggestive that mast cells may be active in human cGVHD. Further studies directly or indirectly addressing mast cells and correlative biomarkers are warranted as these may assist physicians in diagnosing and predicting skin cGVHD as well as expanding the options for treatment.

Our LP/J → C57BL/6J model specifically assesses the role of recipient-derived mast cells. The disease pathology of murine models of cGVHD most often takes place within the first 50 days^43^. However, mast cells can take upwards of 70-80 days to fully reconstitute from the bone marrow to tissue sites^22^. This means that in a theoretical model assessing the role of donor-derived mast cells (C57BL/6J or B6.Cg-Kit^W-sh^/HNihrJaeBsmGlliJ → LP/J), the cells would not have reconstituted from the bone marrow in time to play a role in disease pathogenesis. Furthermore, restoration of mast cells in B6.Cg-Kit^W-sh^/HNihrJaeBsmGlliJ by injection of bone-marrow derived mast cells is not feasible in this model, as injected mast cells do not repopulate the skin after they are injected intravenously or intraperitoneally^22^.

In this current era of targeted therapies, a deeper understanding of disease pathogenesis allows for direct cellular or molecular inhibition. This study is the first *in vivo* evidence of mast cell involvement in dermal cGVHD. Given the mast cell-dependent nature of this model of cGVHD, and the mast cell’s capability to produce many of the same chemokines that we see in the skin of mice with active disease, we propose that mast cell production of chemokines and recruitment of further inflammatory cells may be a critical mechanism of dermal cGVHD pathogenesis. Ibrutinib and ruxolitinib may be capable of modulating this process, and further examination of mast cells in this context may lead to increased therapeutic options for patients who are suffering from cGVHD.

## Acknowledgements

This research was supported by the following core labs: Flow Cytometry and Cell Sorting Shared Resource Facility, Biospecimen Procurement and Translational Pathology Shared Resource Facility, and the Oncogenomics Shared Resource Facility, all through the University of Kentucky Markey Cancer Center (P30CA177558). Research reported in this publication was also supported by an Institutional Development Award (IDeA) from the National Institute of General Medical Sciences of the National Institutes of Health under grant number P20GM103527, and Aperio services were supported through the NIA/NIH ADC P30 AGO28383.

## Materials and methods

### Animals

All animal work was approved by the University of Kentucky Institutional Animal Care and Use Committee and the Department of Laboratory Animal Resources. BALB/cJ, LP/J, C57BL/6J, and B6.Cg-Kit^W-sh^/HNihrJaeBsmGlliJ mice were all purchased from commercially available sources (Stock numbers 000651, 000676, 000664, and 012861, respectively. Jackson Laboratories, Bar Harbor, ME, USA). All mice were female and were used at 8-10 weeks of age.

### Transplant

C57BL/6J or LP/J mice were sacrificed by CO2 inhalation, then bone marrow cells and splenocytes were isolated and counted. C57BL/6J and B6.Cg-Kit^W-sh^/HNihrJaeBsmGlliJ recipients were exposed to a single dose of 850cGy of ionizing radiation from a Cesium source, then injected intravenously with 2.0e6 LP/J splenocytes and 5.0e6 LP/J bone marrow cells. In the syngeneic group, C57BL/6J mice received the same dose of splenocytes and bone marrow cells from a C57BL/6J mouse. After week 7, blood was collected into heparinized tubes (BD Biosciences, San Jose, CA). Heparin tubes were centrifuged at 1500xG for 10 minutes, then plasma was harvested and stored at −80C. Skin, lung, liver, small intestine, colon, kidneys, ear, and tracheal tissue were harvested, trisected, and stored either in formalin for histologic analysis, in RNALater (Millipore-Sigma, St. Louis, MO) for RNA stabilization, or snap-frozen over dry ice for protein analysis, then stored at −80°C. Spleen was harvested for flow cytometric analysis.

### GVHD scoring

Recipient mice were monitored daily for survival, and clinical GVHD was assessed weekly by a lab member not directly involved in the planning of the study. Mice were scored for GVHD by assessment of five clinical parameters: weight loss, posture (hunching), activity, fur texture, and skin integrity. Individual mice from coded cages received a score of 0 to 2 for each criterion (maximum score of 10) that was used as an index of severity and progression of disease^16^.

### Mast cell culture

Bone marrow was extracted from femurs and tibias isolated from BALB/cJ or C57BL/6J mice and cultured in DMEM media (Lonza, Basel, Switzerland) with 10% heat-inactivated fetal bovine serum (Corning Inc., Corning, NY) and 1% penicillin/streptomycin (Quality Biological, Gaithersburg, MD) at 37C, 5% CO2, in a humidified incubator. After 24 hours, the suspension cells were removed and culture in a new flask along with 10ng/ml recombinant murine IL-3 (Peprotech, Rocky Hill, NJ). Cells were passaged twice per week, each time leaving behind any adherent cells. After 4-6 weeks, cell phenotype was confirmed by visual morphology and flow cytometry.

### Mast cell irradiation and viability assay

Viable mast cells were counted by hemacytometer via trypan blue exclusion and plated at 1e6/ml, then exposed to 8.5 or 20Gy single-dose radiation from a cesium source. Cells were approximately 99% viable prior to irradiation. Every 24 hours, cells were counted and assessed for viability by trypan blue exclusion out to 96 hours post-irradiation.

### Mast cell stimulation and inhibition assays

Mast cells were incubated overnight with 100ng/ml murine monoclonal anti-DNP IgE, clone SPE-7 (Millipore-Sigma, St. Louis, MO) at 37C. The next day, cells were centrifuged and washed twice with fresh media to remove unbound IgE and cells were replated in fresh media with 10ng/ml IL-3. Stimulation was to the relevant wells added in the form of 10ng/ml IL-33 and/or 100ng/ml DNP-BSA for 24 hours at 37C, then cells were centrifuged and supernatant was collected. For the drug inhibition assays, ibrutinib, ruxolitinib, imatinib, and fingolimod were purchased from Cayman Chemical (Ann Arbor, MI) and reconstituted per the manufacturer’s instructions. Drug stocks were diluted in media and incubated with mast cells after the overnight IgE incubation step. Cells were treated for one hour at 37C, then washed out with fresh media twice. Cells were then stimulated as per above.

### Degranulation by CD107a expression

Cells were incubated at 1e6/ml with IgE overnight then washed and plated in new media, as above. 0.66ul/ml Golgistop (BD Biosciences, San Jose, CA) was added along with AF594-CD107a antibody, clone 1D4B, (Biolegend, San Diego, CA), then cells were stimulated with 100ng/ml DNP-BSA and/or 50ng/ml SCF (Peprotech) for 4 hours at 37C. Cells were then collected, washed, and fixed/permeabilized with Fix/Perm reagent (BD Biosciences). Cells were then stained with BV421-CD117 (clone 2B8) and PE-FCeR1a (clone MAR-1) (Biolegend) in perm/wash buffer for 20 minutes at 4C, washed, and analyzed by flow cytometry for CD117/FCeR1a +/+ CD107 +/-expression.

### Histology

Organs were collected from mice and suspended in 4% formalin for 48 hours, then 70% ethanol for 7 days. After this, samples were cut into slices, processed, and embedded in paraffin blocks. All sections shown were sectioned at 5 micron thickness. Masson’s trichrome staining was performed according to manufacturer protocol (Electron Microscopy Sciences, Hatfield, PA). H+E stains were performed by the University of Kentucky Surgical Pathology core and assessed by a blinded, board-certified pathologist according to a previously published organ-specific GVHD scoring system^16,44^. Skin slides were stained with Toluidine Blue O (Millipore Sigma) and assessed as toluidine blue-positive cells per high-power field. Counts were verified by a separate, blinded researcher.

### Immunohistochemistry

Slides were baked at 80C for 15 minutes, then rehydrated through xylene, 100%, 95%, and 70% ethanol into H2O. Antigen retrieval was performed in citrate Target Retrieval Solution (Agilent, Santa Clara, CA), pH 6.0, in a pressure cooker. Slides were blocked for 30 minutes in PowerBlock (Biogenex, Fremont, CA), then incubated with primary antibody at a 1:200 dilution in PowerBlock. Antibodies used were rabbit anti-mouse CD3e, clone D4V8L (Cell Signaling Technologies, Danvers, MA), and mouse anti-human tryptase, clone AA1 (Abcam, Boston, MA). Slides were washed in PBS + 0.1% Tween 20 (PBS-T) and then incubated with secondary antibody. For CD3 staining, a drop of Dako Envision+ HRP Anti-Rabbit Polymer (Agilent) was added to slides for one hour, then washed with PBS-T. For tryptase staining, Goat Anti-Mouse Immunoglobulin/HRP (Agilent) was used at a 1:1 dilution for 10 minutes and then washed with PBS-T. After washing, DAB chromogen (Abcam) was prepared according to manufacturer’s instructions and added to slides while observing the staining under a microscope for 30 seconds (tryptase staining) or 2 minutes (CD3 staining). The reaction was quenched in diH2O, then dehydrated to xylene. Slides were mounted using Richard-Allan Scientific mounting medium (Thermo Fisher Scientific, Waltham, MA).

### Microscopy and analysis

Slides were imaged by Aperio Scanscope XT digital slide scanner or by a color camera system attached to a Nikon Eclipse 80i microscope. All scale bars are 100um. Trichrome and CD3 IHC slides were digitally scanned by an Aperio ScanScope XT system. Dermal thickness was measured with the ruler tool across 10-20 areas of tissue. Aperio methodology and results were checked and verified by CDJ. CD3 and tryptase positivity was measured across the whole tissue by the Aperio Positive Pixel Count V9 algorithm. Counts were also verified by ImageJ IHC profiler analysis by a blinded, independent researcher, RK.

### Flow cytometry

Splenocytes were phenotyped using two different antibody panels. The first panel consisted of lymphocyte and T-cell stains and allowed for identification of CD45, CD3, CD4, CD8, and FoxP3 subsets. The second panel identified B-cells, neutrophils, macrophages, and dendritic cells using stains for CD19, CD11c, CD11b, F4/80, and Ly6G (Gating scheme for both panels in Supplementary Fig. 5). Splenic tissue was forced through a 70 micron filter to create a single cell suspension and cells were washed with PBS containing 4% FBS and incubated with FcR-block anti-mouse CD16/CD32 (Clone 93, eBioscience, San Diego, CA). Cells then were incubated in a pre-optimized concentration of antibodies in a total volume of 100μL. For intracellular staining for FoxP3, fixation and permeabilization were done as per protocol using the fixation/permeabilization concentrate and diluent (eBioscience). The following antibody clones were used for splenocyte characterization:

FITC-CD3 (REA641), APC-CD4 (GK1.5), APC-Vio770-CD45 (30F11), APC-CD11c (N418), PE-CD19 (6D5), FITC-F4/80 (REA126) (Miltenyi Biotec, Bergisch Gladbach, Germany)

PerCP-Cy5.5-CD11b (M1/70), PerCP-Cy5.5-CD8 (53-6.7) (Biolegend, San Jose, CA)

PE-Cy7-Ly6G (RB6-8C5), APC-eFluor780-MHCII (M5/114.15.2), PE-FoxP3 (FJK-16s) (eBioscience)

Mast cells were phenotyped using the following antibody clones:

BV421-CD117 (2B8), PE-FCeR1a (MAR-1) (Biolegend)

All cells were analyzed on a BD LSR II cytometer (BD Biosciences) using single-stained OneComp eBeads (Thermo Fisher Scientific, Waltham, MA) to determine compensation values. All gates were set using fluorescence-minus-one (FMO) controls. Flow cytometry data were analyzed using FlowJo V10 software (FlowJo Inc., Ashland, OR).

### RNA isolation and NanoString analysis

Tissue was frozen in RNALater as described in “Transplant” methods. The tissue was thawed, then suspended in 1ml of Ribozol RNA isolation solution (VWR, Radnor, PA), and homogenized using a Tissuemiser homogenizer (Thermo Fisher, Waltham, MA). RNA was isolated from the homogenate per the Ribozol manufacturer’s instructions. RNA was quantitated using the Qubit RNA BR assay kit in the Qubit Fluorometer (Thermo Fisher). 100ng of RNA was loaded into the cartridge of the NanoString Myeloid Innate Immunity panel and run on the nCounter Flex system at the University of Kentucky Clinical Genomics core per the manufacturer’s instructions (NanoString Technologies, Seattle, WA). Data normalization and analysis was performed in the nSolver software.

### Real-time quantitative PCR

For cDNA synthesis and qPCR, RNA was quantified using a Nanodrop instrument (Thermo Fisher), and 100ng was loaded into the iScript cDNA Synthesis Kit (Bio-Rad, Hercules, CA). For IL21 gene analysis, a predesigned primer probe mix was purchased from Integrated DNA Technologies (Assay ID: Mm.PT.58.7853071, Integrated DNA Technologies, San Jose, CA) and normalized using ACTB as a housekeeping gene (Assay ID: Mm.PT.39a.22214843.g). cDNA was assayed using the Luna Universal Probe qPCR master-mix, (New England Biolabs, Ipswich, MA) on an Applied Biosystems StepOne Real-Time PCR System (Applied Biosystems, Foster City, CA). Relative quantitation was performed using the delta-delta CT method, described previously^45^.

### Tissue lysate preparation

Tissue was snap-frozen on dry ice as described in the “Transplant” methods section. The tissue was thawed, then suspended in 500ml 1x RIPA buffer (Thermo Fisher, Waltham, MA) with protease inhibitors added (cOmplete Protease Inhibitor Cocktail, Roche, Basel, Switzerland). Tissue was homogenized using a Tissuemiser homogenizer (Thermo Fisher). Homogenate was vortexed for 30 seconds, then put on ice for 10 minutes. This procedure was repeated 3 times, then homogenates were spun down at 14,000xG for 10 minutes at 4C. Supernatant was moved to a new tube and quantified by Pierce BCA protein assay kit (Thermo Fisher).

### Measurement of cytokines and chemokines

The levels of TNF-α, IL-6, IFN-γ, IL-2, IL-4, IL-10 and IL-17α were determined in plasma samples through the Mouse Th1/Th2/Th17 Cytometric Bead Array kit (BD Biosciences, San Jose, CA). Bead fluorescence was measured using the BD Biosciences LSRII flow cytometer and analyzed with the accompanying software. Protein levels of CCL5, CCL11, CCL17, CXCL1, CXCL9, CXCL10, CXCL13, CXCL5, CCL22, CCL2, CCL3, and CCL4 in tissue and mast cell supernatants were analyzed with the LEGENDplex Mouse Proinflammatory Chemokine Panel per the manufacturer’s instructions. Supernatants were loaded without dilution. Approximately 100 micrograms of tissue lysate were loaded for analysis of chemokine levels in the skin, although final values displayed in the paper were normalized to picograms per milliliter per microgram of protein loaded. Bead fluorescence was measured using the BD Biosciences LSRII flow cytometer and analyzed in the Biolegend LEGENDplex data analysis software.

### Fibroblast proliferation assay

Fibroblasts were cultured from the skin of C57BL/6J mice as previously described^46^. After 3 weeks of culture, cells were nearly confluent in a 10cm petri dish. Media was aspirated off and 3 milliliters of trypsin was added to the dish for 5 minutes. After 5 minutes, cells were scraped off using a sterile scraper and spun down at 350xG for 5 minutes. Cells were resuspended in 6 milliliters and plated equally into a 6 well plate in DMEM + 10% FBS + 1% penicillin/streptomycin. After 4 hours to allow the cells to adhere, cells were counted on an inverted microscope across 4 fields of view per well at 200x magnification. At this point, 1ml of mast cell supernatants or fresh media was added to the 1ml of culture in each well. Cells were counted again 24 hours later, and again after 48 hours.

### Human tissue samples

All samples were from patients at the University of Kentucky Markey Cancer Center who had undergone an allo-HSCT with a subsequent diagnosis of chronic GVHD. Chart review and sample acquisition was performed by the honest broker of the University of Kentucky Biospecimen Procurement and Translational Pathology Core and samples were sectioned from paraffin-embedded tissue. All samples were deidentified and were not able to be linked to patient records by any investigators involved in the study. As a result of this, the University of Kentucky Institutional Review Board deemed that these samples did not meet the Department of Health and Human Services (DHHS) definition of human subjects or the Food & Drug Administration’s (FDA) definition of human subjects.

### Data sharing and availability

Nanostring data is stored in the publicly available NCBI Gene Expression Omnibus database (accession number GSE128704). Other data in this study is available from the corresponding author upon request.

### Figure preparation and statistics

All figures were prepared in GraphPad Prism 6 and 7 (GraphPad, San Diego, CA), FlowJo V10 (FlowJo Inc., Ashland, OR), or NanoString nSolver (NanoString Technologies, Seattle, WA) software packages. Experimental data are expressed as means ± standard deviation unless noted otherwise in the figure legend. Differences between two groups were analyzed by the two-tailed independent t-test. Differences between three or more groups were analyzed by one-way ANOVA, with a Tukey’s post-hoc multiple comparisons test to determine the significance between groups. P values ≤0.05 were considered statistically significant. P value 0.01 to 0.05 = *, P value 0.001 to 0.01 = **, P value 0.0001 to 0.001 = ***, P value <0.0001 = ****, NS = not significant

---

**Supplementary Figure 1:**
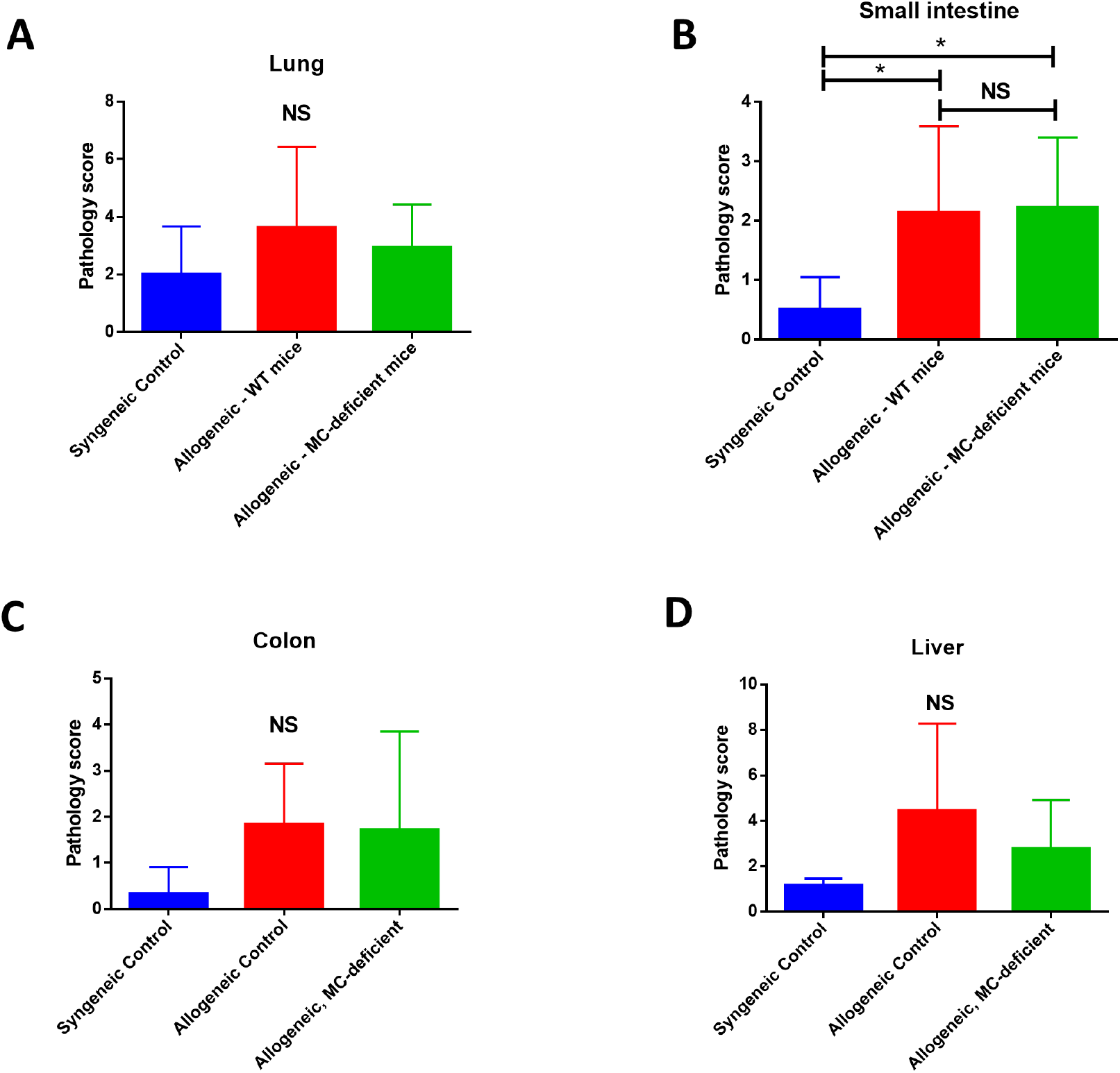
Pathology is not significantly different after allogeneic transplant between allo-WT and allo-MCd in lung, small intestine, colon, or liver. Pathology score is unchanged between all groups in A) lung, B) small intestine, C) colon, or D) liver. Scoring was performed as described previously^16,44^ by a blinded pathologist. Syngeneic (n=6), allo-WT (n=16), and allo-MCd (n=14). Data is combined from two independent transplants. P value 0.01 to 0.05 = *, P value 0.001 to 0.01 = **, P value 0.0001 to 0.001 = ***, P value <0.0001 = ****, NS = not significant

**Supplementary Figure 2:**
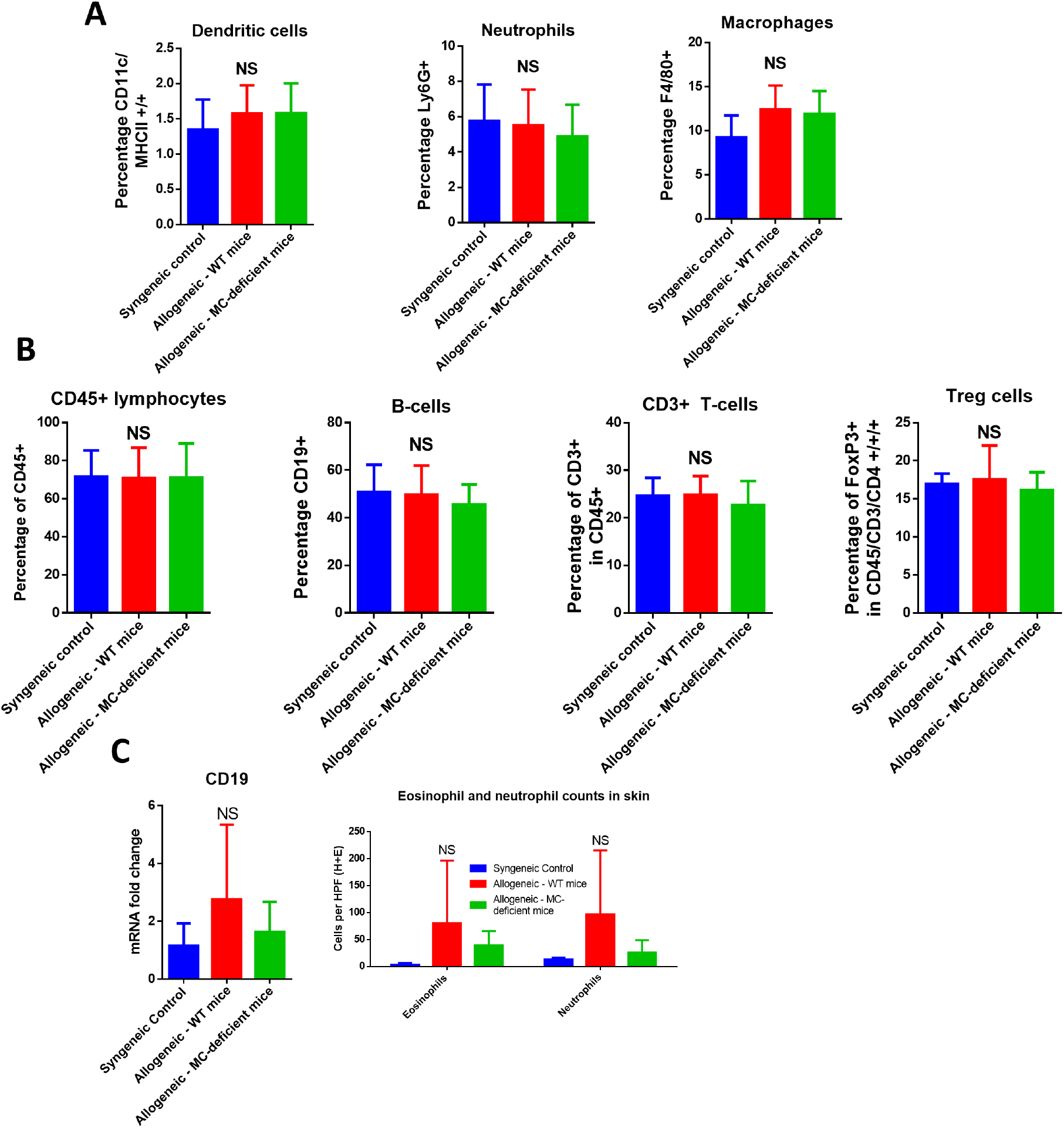
Markers of many immune subsets in the spleen and skin are not significantly changed. A) Myeloid subsets are unchanged in the spleen 7 weeks after allogeneic transplant. MHCII/CD11c +/+ dendritic cells, Ly6G+ neutrophils, or CD11b/F4/80 +/+ macrophages have no significant differences in proportion or overall count (data not shown) in the spleen after induction of cGVHD. B) There were no significant differences in splenic proportion or count (data not shown) of the lymphoid subsets analyzed (CD45+ lymphocytes, CD45/CD19 +/+ B-cells, CD45/CD3 +/+ T-cells, CD45/CD3/CD4/FoxP3 +/+/+/+ T-regulatory cells). This implies that the dermal cGVHD symptomology evident in these mice is driven more strongly by local factors than purely by increased alloreactivity, a conclusion which is consistent with many theories regarding the pathogenesis of fibrotic cGVHD. C) There is no significant difference in the skin in CD19 transcript (measured by qPCR) or eosinophil/neutrophil counts (counted by a pathologist by H+E morphology). P value 0.01 to 0.05 = *, P value 0.001 to 0.01 = **, P value 0.0001 to 0.001 = ***, P value <0.0001 = ****, NS = not significant

**Supplementary Figure 3:**
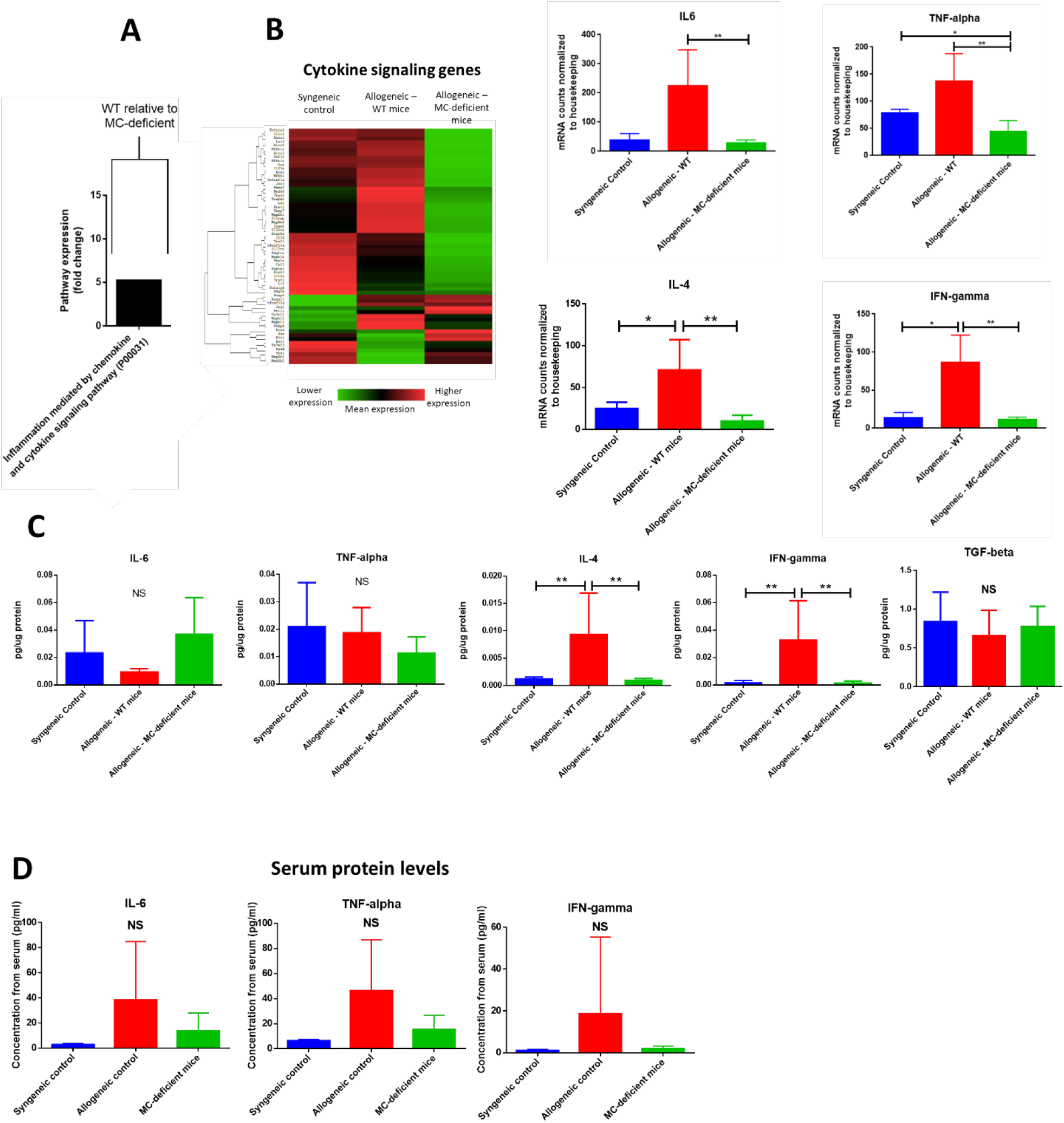
Pathogenic cytokines are expressed at low levels in the skin and are largely unchanged between groups. A) PANTHER pathway analysis demonstrating an increase in genes related to “Inflammation mediated by chemokine and cytokine signaling” in allo-WT relative to allo-MCd. B) Heatmap analysis and selected genes showing lowered expression of cytokine signaling genes in allo-MCd animals compared to allo-WT animals as measured by NanoString. Heatmaps and gene pathway annotations were generated using NanoString nSolver software. C) Protein levels were measured in the skin for IL-6, TNF-alpha, IL-4, and IFN-gamma. D) Protein levels in plasma (syngeneic n=3, allo-WT n=8, and allo-MCd n=7) P value 0.01 to 0.05 = *, P value 0.001 to 0.01 = **, P value 0.0001 to 0.001 = ***, P value <0.0001 = ****, NS = not significant

**Supplementary Figure 4:**
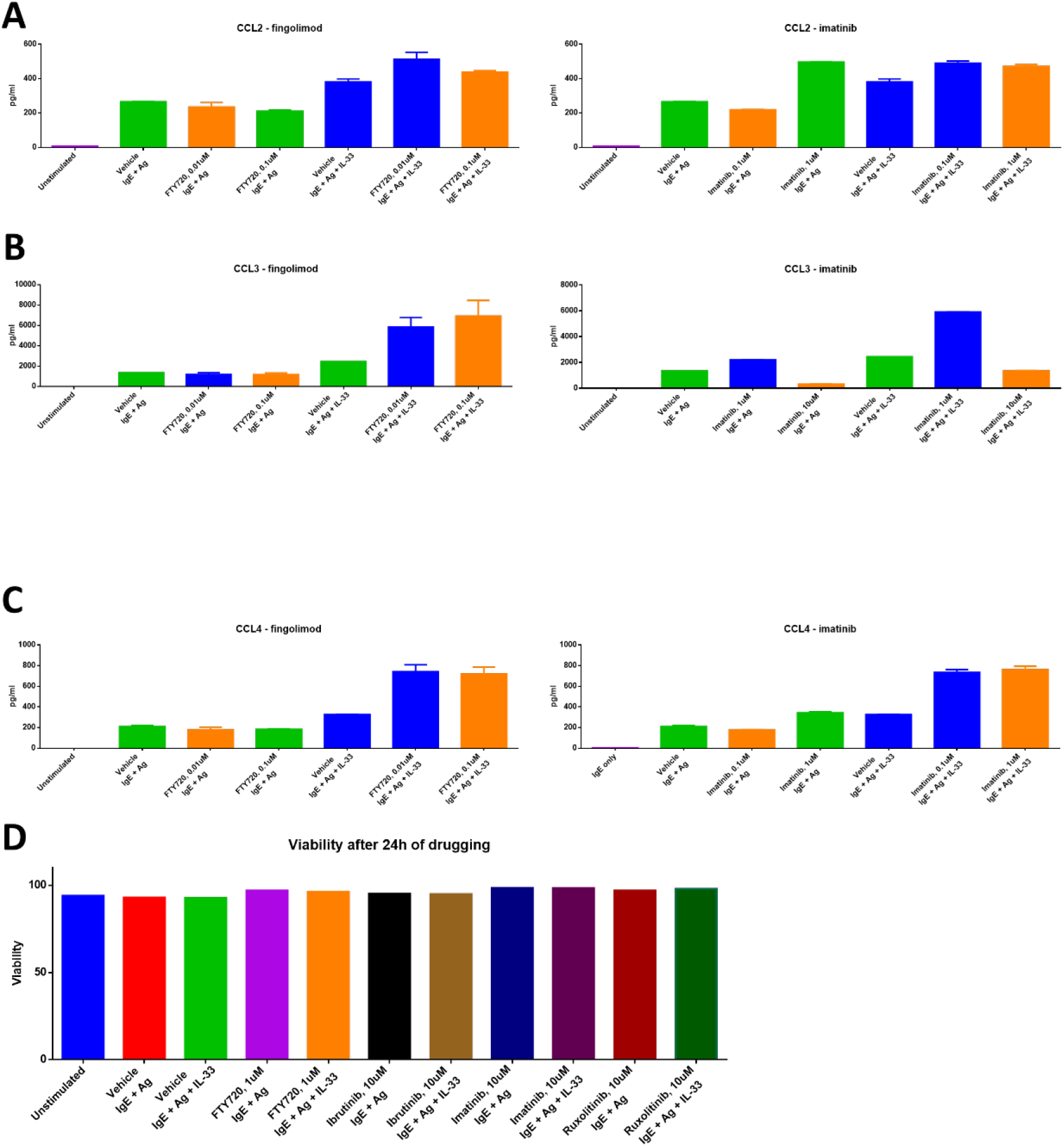
Chemokine production is not reduced after treatment with imatinib or fingolimod and cell viability is unaffected by drugging. Mast cells produce high levels of A) CCL2, B) CCL3, and C) CCL4 upon stimulation with IgE + antigen or IgE + antigen + IL-33 (column 1 vs. columns 2 and 6). Production of these chemokines is not decreased by treatment with either imatinib or fingolimod. Results shown are representative of 2-4 independent assays. Error bars are the SD of technical replicates. Chemokine assays were performed using the LEGENDplex Inflammatory Chemokine Assay kit, which measures levels of 13 chemokines. Mast cells did not produce significant amounts of CCL5, CCL11, CCL17, CXCL1, CXCL9, CXCL10, CXCL13, CXCL5, or CCL22 (data not shown). D) Mast cell viability was unaffected after 24 hours of drugging with either imatinib, fingolimod, ibrutinib, or ruxolitinib. P value 0.01 to 0.05 = *, P value 0.001 to 0.01 = **, P value 0.0001 to 0.001 = ***, P value <0.0001 = ****, NS = not significant

**Supplementary Figure 5:**
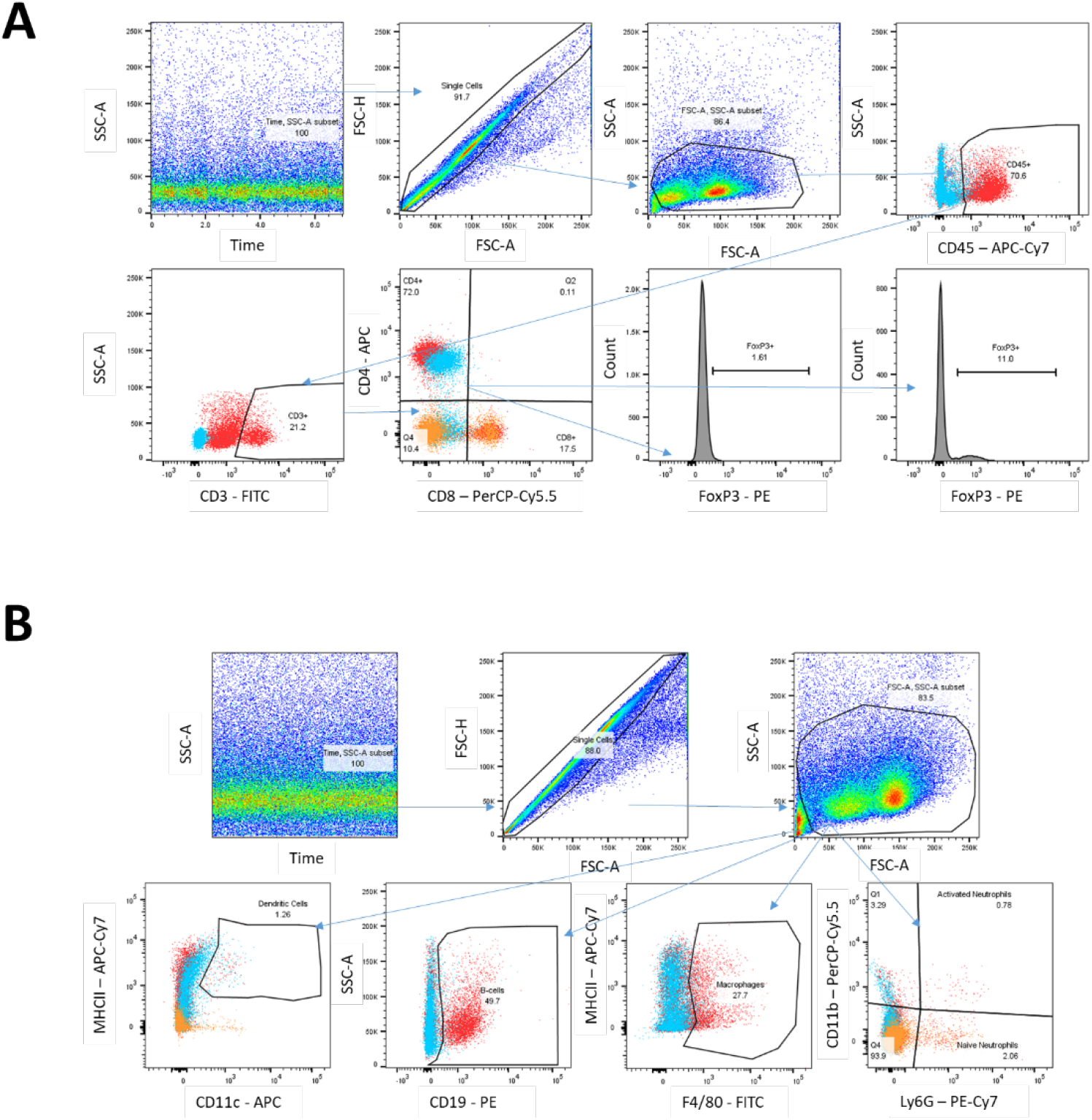
Flow cytometry gating schemes. Gating schema for flow cytometry panels run on spleen (Supp. Fig. 2). A) Gating scheme for a panel to assay T-cell subsets in the spleen. B) Gating scheme for a panel to assay myeloid subsets and B-cells in the spleen. Red samples are fully stained, while blue or orange are FMO controls.

## References

1. Copelan, E. A. Hematopoietic Stem-Cell Transplantation. N. Engl. J. Med. 354, 1813–1826 (2006).

2. Arai, S. et al. Increasing Incidence of Chronic Graft-versus-Host Disease in Allogeneic Transplantation – A Report from CIBMTR. Biol. Blood Marrow Transplant. J. Am. Soc. Blood Marrow Transplant. 21, 266–274 (2015).

3. Flowers, M. E.D. & Martin, P. J. How we treat chronic graft-versus-host disease. Blood 125, 606–615 (2015).

4. Galli, S. J. & Tsai, M. Mast cells in allergy and infection: Versatile effector and regulatory cells in innate and acquired immunity. Eur. J. Immunol. 40, 1843–1851 (2010).

5. Pincha, N. et al. PAI1 mediates fibroblast–mast cell interactions in skin fibrosis. J. Clin. Invest. 128, 1807–1819 (2018).

6. Hügle, T. Beyond allergy: the role of mast cells in fibrosis. Swiss Med. Wkly. 144, w13999 (2014).

7. Veerappan, A. et al. Mast cells: a pivotal role in pulmonary fibrosis. DNA Cell Biol. 32, 206–218 (2013).

8. Claman, H. N., Choi, K. L., Sujansky, W. & Vatter, A. E. Mast cell ‘disappearance’ in chronic murine graft-vs-host disease (GVHD)-ultrastructural demonstration of ‘phantom mast cells’. J. Immunol. 137, 2009–2013 (1986).

9. Claman, H. N. Mast Cell Depletion in Murine Chronic Graft-versus-Host Disease. J. Invest. Dermatol. 84, 246–248 (1985).

10. Levi-Schaffer, F., Mekori, Y. A., Segal, V. & Claman, H. N. Histamine release from mouse and rat mast cells cultured with supernatants from chronic murine graft-vs-host splenocytes. Cell. Immunol. 127, 146–158 (1990).

11. Claman, H. N. Mast cells, T cells and abnormal fibrosis. Immunol. Today 6, 192–195 (1985).

12. Levi-Schaffer, F., Segal, V., Baram, D. & Mekori, Y. A. Effect of coculture of rodent mast cells with murine chronic graft-versus-host disease (cGVHD)-derived fibroblasts. J. Allergy Clin. Immunol. 89, 501–509 (1992).

13. Grimbaldeston, M. A. et al. Mast Cell-Deficient W-sash c-kit Mutant KitW-sh/W-sh Mice as a Model for Investigating Mast Cell Biology in Vivo. Am. J. Pathol. 167, 835–848 (2005).

14. Soule, B. P. et al. Effects of Gamma Radiation on FcεRI and TLR-Mediated Mast Cell Activation. J. Immunol. 179, 3276–3286 (2007).

15. Hamilton, B. L. & Parkman, R. Acute and chronic graft-versus-host disease induced by minor histocompatibility antigens in mice. Transplantation 36, 150–155 (1983).

16. Cooke, K. R. et al. An experimental model of idiopathic pneumonia syndrome after bone marrow transplantation: I. The roles of minor H antigens and endotoxin. Blood 88, 3230–3239 (1996).

17. Martires, K. J. et al. Sclerotic-type chronic GVHD of the skin: clinical risk factors, laboratory markers, and burden of disease. Blood 118, 4250–4257 (2011).

18. Shulman, H. M. et al. HISTOPATHOLOGIC DIAGNOSIS OF CHRONIC GRAFT VERSUS HOST DISEASE. Biol. Blood Marrow Transplant. J. Am. Soc. Blood Marrow Transplant. 21, 589–603 (2015).

19. Taylor, D. K. et al. T follicular helper-like cells contribute to skin fibrosis. Sci. Transl. Med. 10, (2018).

20. Pohlers, D. et al. TGF-β and fibrosis in different organs — molecular pathway imprints. Biochim. Biophys. Acta BBA - Mol. Basis Dis. 1792, 746–756 (2009).

21. Wynn, T. A. Cellular and molecular mechanisms of fibrosis. J. Pathol. 214, 199–210

22. Wolters, P. J. et al. Tissue-selective mast cell reconstitution and differential lung gene expression in mast cell-deficient Kit(W-sh)/Kit(W-sh) sash mice. Clin. Exp. Allergy J. Br. Soc. Allergy Clin. Immunol. 35, 82–88 (2005).

23. Mi, H. et al. PANTHER version 11: expanded annotation data from Gene Ontology and Reactome pathways, and data analysis tool enhancements. Nucleic Acids Res. 45, D183–D189 (2017).

24. Luther, S. A. & Cyster, J. G. Chemokines as regulators of T cell differentiation. Nat. Immunol. 2, 102–107 (2001).

25. Wang, A., Patterson, S., Marwaha, A., Tan, R. & Levings, M. CCL3 and CCL4 secretion by T regulatory cells attracts CD4+ and CD8+ T cells (P1077). J. Immunol. 190, 121.10–121.10 (2013).

26. Mukai, K., Tsai, M., Saito, H. & Galli, S. J. Mast cells as sources of cytokines, chemokines and growth factors. Immunol. Rev. 282, 121–150 (2018).

27. Choi, S. W. et al. CCR1/CCL5 (RANTES) receptor-ligand interactions modulate allogeneic T-cell responses and graft-versus-host disease following stem-cell transplantation. Blood 110, 3447–3455 (2007).

28. Blirando, K. et al. Mast Cells Are an Essential Component of Human Radiation Proctitis and Contribute to Experimental Colorectal Damage in Mice. Am. J. Pathol. 178, 640–651 (2011).

29. Qu, Z. et al. Mast cells are a major source of basic fibroblast growth factor in chronic inflammation and cutaneous hemangioma. Am. J. Pathol. 147, 564–573 (1995).

30. Leveson-Gower, D. B. et al. Mast cells suppress murine GVHD in a mechanism independent of CD4+ CD25+ regulatory T cells. Blood 122, 3659–3665 (2013).

31. Blazar, B., White, E. S. & Couriel, D. Understanding Chronic GVHD from Different Angles. Biol. Blood Marrow Transplant. J. Am. Soc. Blood Marrow Transplant. 18, S184–S188 (2012).

32. Abel, M. & Vliagoftis, H. Mast Cell-Fibroblast Interactions Induce Matrix Metalloproteinase-9 Release from Fibroblasts: Role for IgE-Mediated Mast Cell Activation. J. Immunol. 180, 3543–3550 (2008).

33. Arbi, S., Eksteen, E. C., Oberholzer, H. M., Taute, H. & Bester, M. J. Premature collagen fibril formation, fibroblast-mast cell interactions and mast cell-mediated phagocytosis of collagen in keloids. Ultrastruct. Pathol. 39, 95–103 (2015).

34. Garbuzenko, E. et al. Human mast cells stimulate fibroblast proliferation, collagen synthesis and lattice contraction: a direct role for mast cells in skin fibrosis. Clin. Exp. Allergy 32, 237–246 (2002).

35. Hellman, L. T., Akula, S., Thorpe, M. & Fu, Z. Tracing the Origins of IgE, Mast Cells, and Allergies by Studies of Wild Animals. Front. Immunol. 8, (2017).

36. Dubovsky, J. A. et al. Ibrutinib treatment ameliorates murine chronic graft-versus-host disease. J. Clin. Invest. 124, 4867–4876 (2014).

37. Miklos, D. et al. Ibrutinib for chronic graft-versus-host disease after failure of prior therapy. Blood 130, 2243–2250 (2017).

38. Sarmiento Maldonado, M. et al. Compassionate use of ruxolitinib in acute and chronic graft versus host disease refractory both to corticosteroids and extracorporeal photopheresis. Exp. Hematol. Oncol. 6, 32 (2017).

39. Study of Ruxolitinib in Sclerotic Chronic Graft-Versus-Host Disease After Failure of Systemic Glucocorticoids - Full Text View - ClinicalTrials.gov. Available at: https://clinicaltrials.gov/ct2/show/NCT03616184. (Accessed: 14th March 2019)

40. A Study of Ruxolitinib vs Best Available Therapy (BAT) in Patients With Steroid-refractory Chronic Graft vs. Host Disease (GvHD) After Bone Marrow Transplantation (REACH3) - Full Text View - ClinicalTrials.gov. Available at: https://clinicaltrials.gov/ct2/show/NCT03112603. (Accessed: 14th March 2019)

41. Morales, J., Falanga, Y., Depcrynski, A., Fernando, J. & Ryan, J. Mast cell homeostasis and the JAK–STAT pathway. Genes Immun. 11, 599–608 (2010).

42. Huang, W., Morales, J. L. & August, A. Itk and Btk regulate mast cell responses to lipopolysaccharide: firing or dampening? (151.18). J. Immunol. 186, 151.18–151.18 (2011).

43. Schroeder, M. A. & DiPersio, J. F. Mouse models of graft-versus-host disease: advances and limitations. Dis. Model. Mech. 4, 318–333 (2011).

44. Kaplan, D. H. et al. Target Antigens Determine Graft-versus-Host Disease Phenotype. J. Immunol. 173, 5467–5475 (2004).

45. Fleige, S. et al. Comparison of relative mRNA quantification models and the impact of RNA integrity in quantitative real-time RT-PCR. Biotechnol. Lett. 28, 1601–1613 (2006).

46. Seluanov, A., Vaidya, A. & Gorbunova, V. Establishing Primary Adult Fibroblast Cultures From Rodents. J. Vis. Exp. JoVE (2010). doi:10.3791/2033

