## Supplementary figures for "Mast cells are mediators of fibrosis and effector cell recruitment in dermal chronic graft-versus-host disease"

Supplementary 1– Pathology is not significantly different after allogeneic transplant between allo-WT and allo-MCd in lung, small intestine, colon, or liver

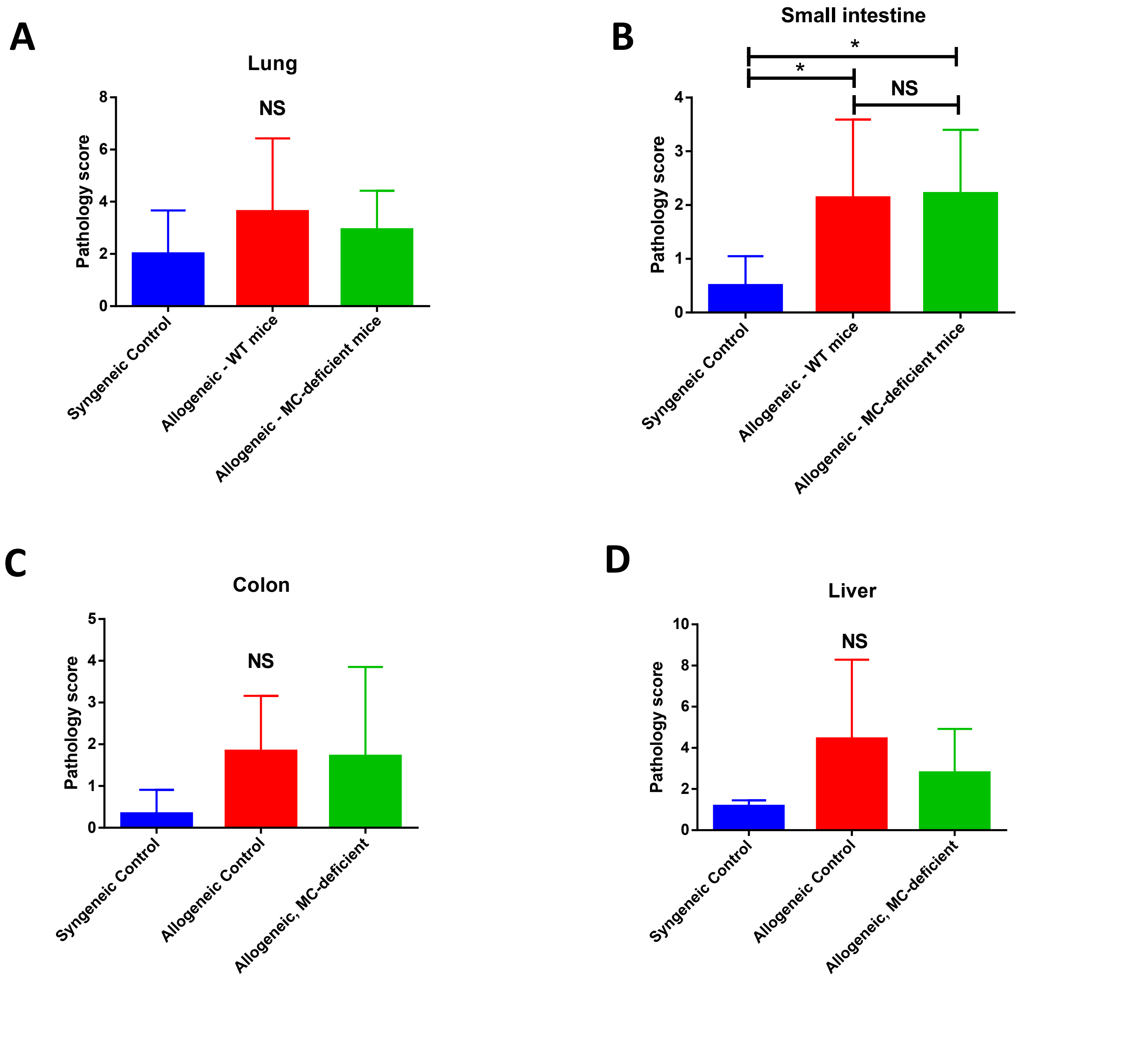

**Supplementary 2 –** Markers of many immune subsets in the spleen and skin are not significantly changed.

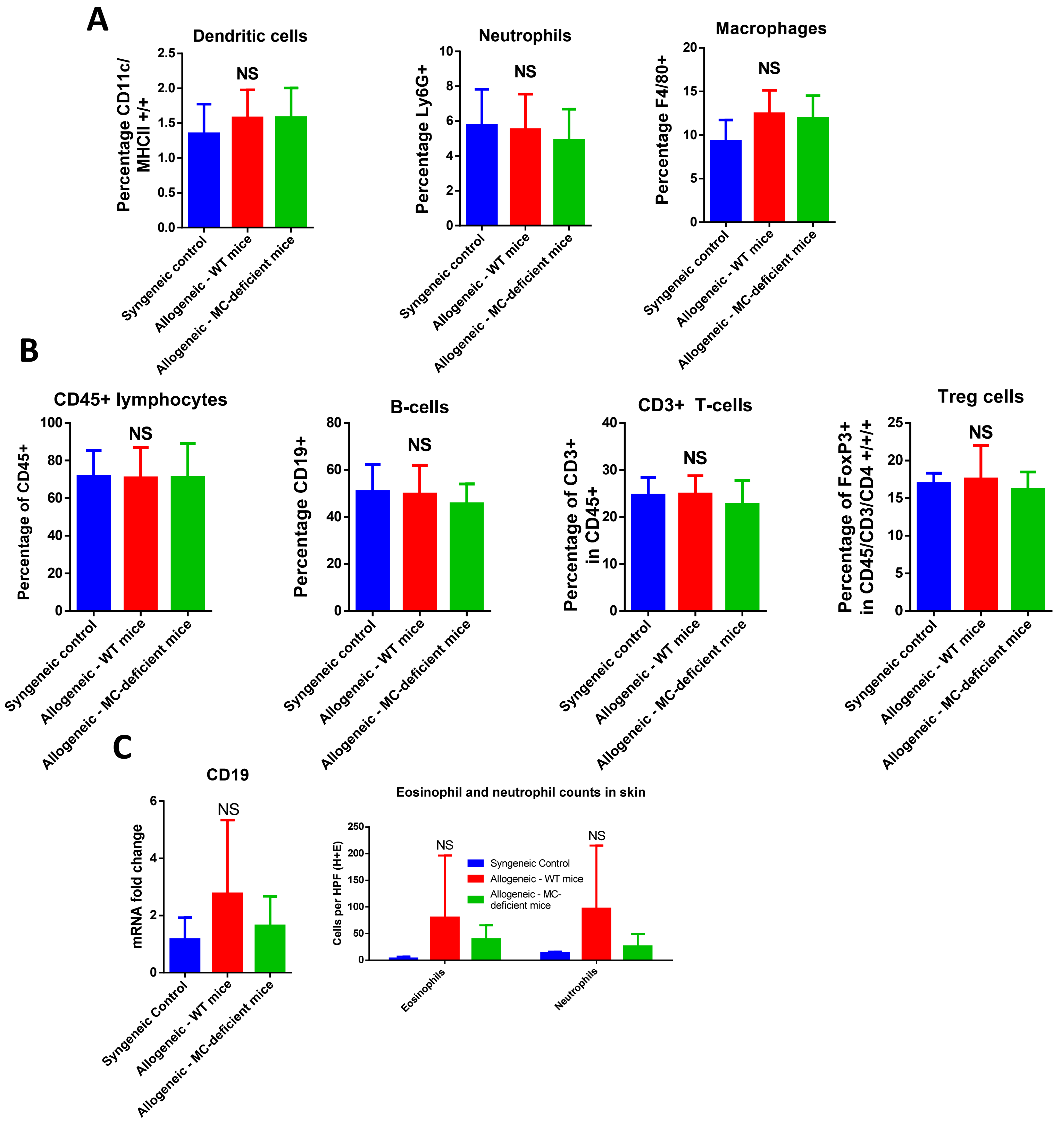

### Supplementary 3 – Pathogenic cytokines are expressed at low levels in the skin and are largely unchanged between groups

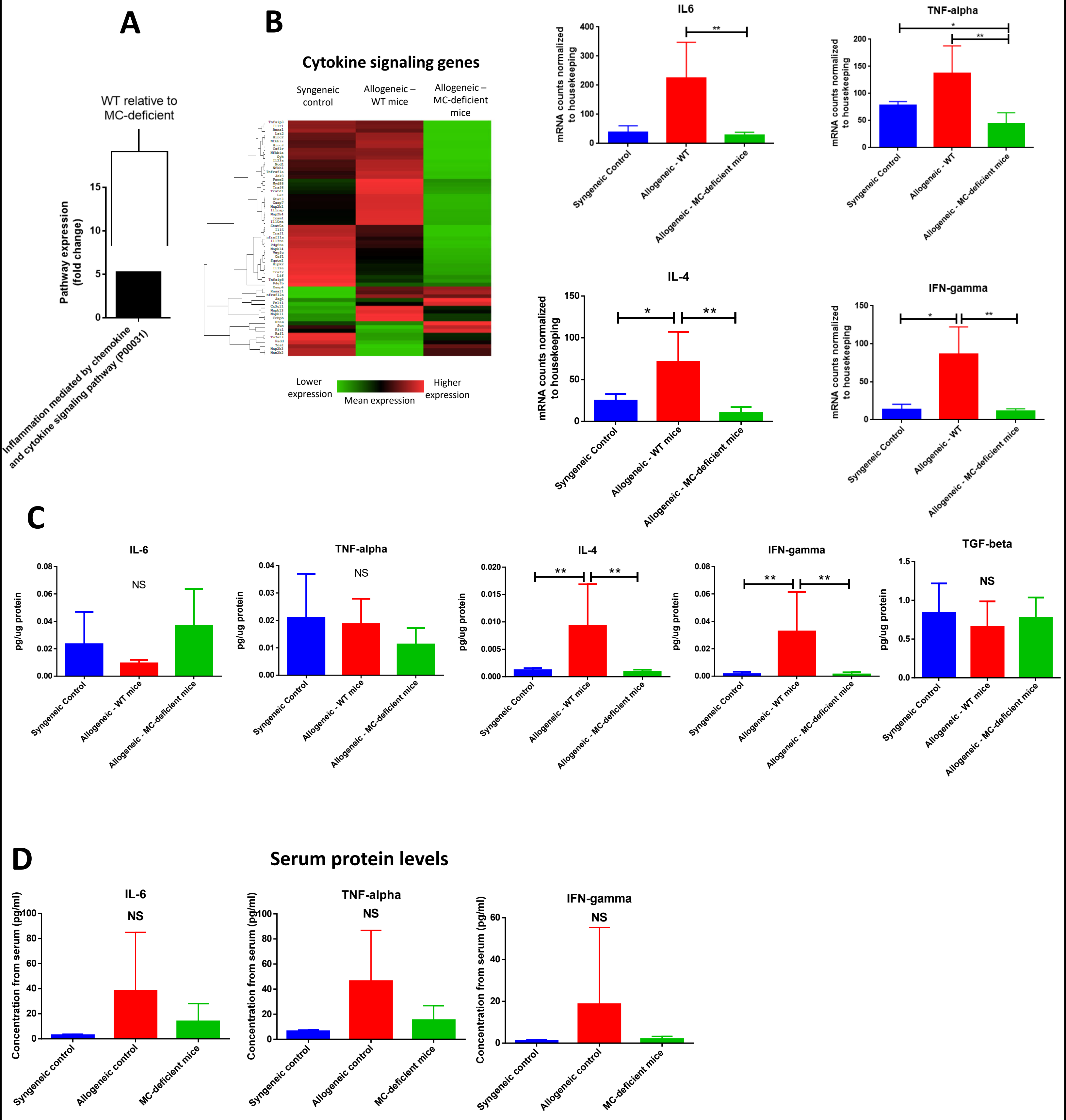

**Supplementary Figure 4:** Chemokine production is not reduced after treatment with imatinib or fingolimod and cell viability is unaffected by drugging

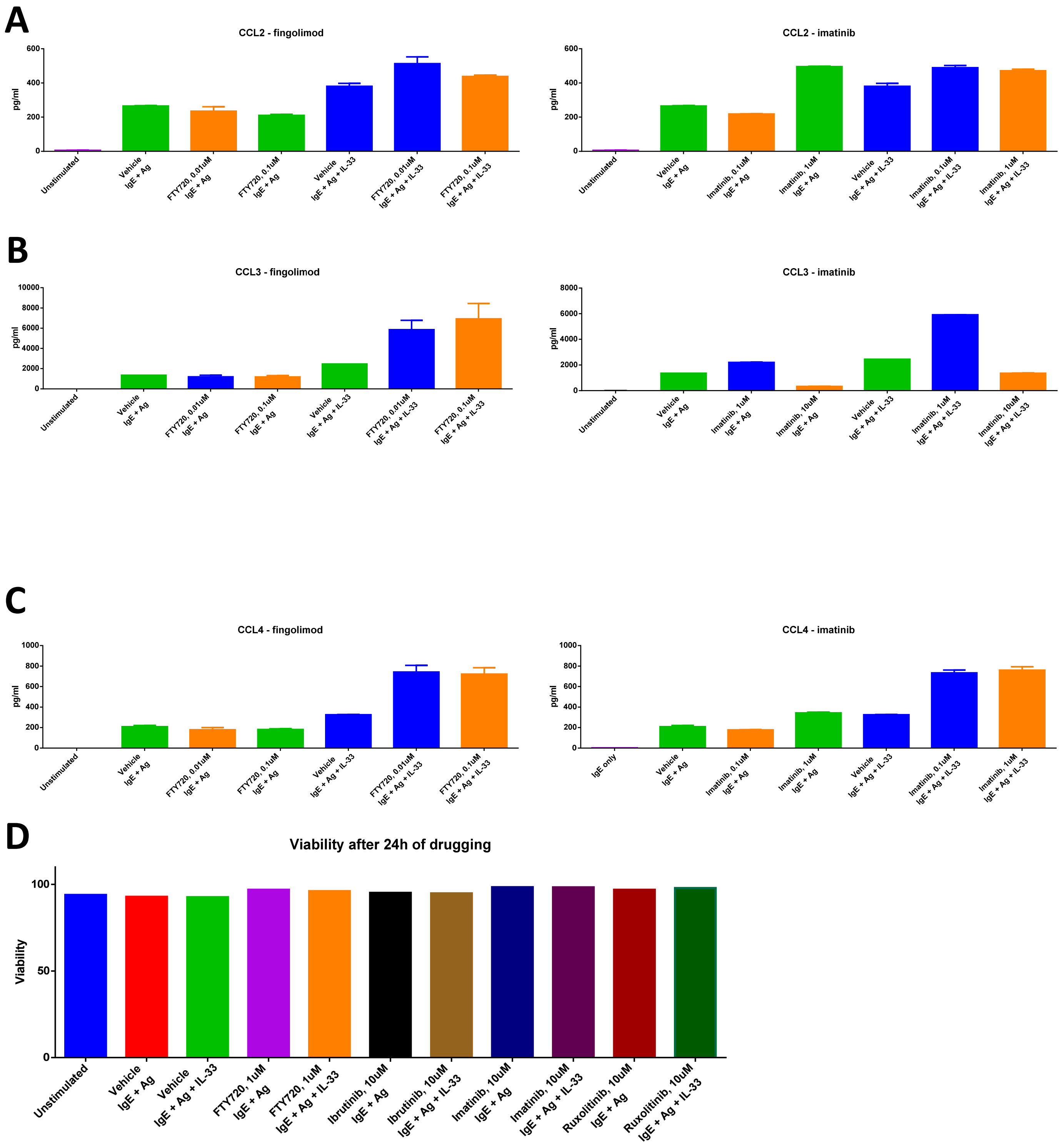

### Supplementary 5– Gating schemes for flow cytometry

**A**

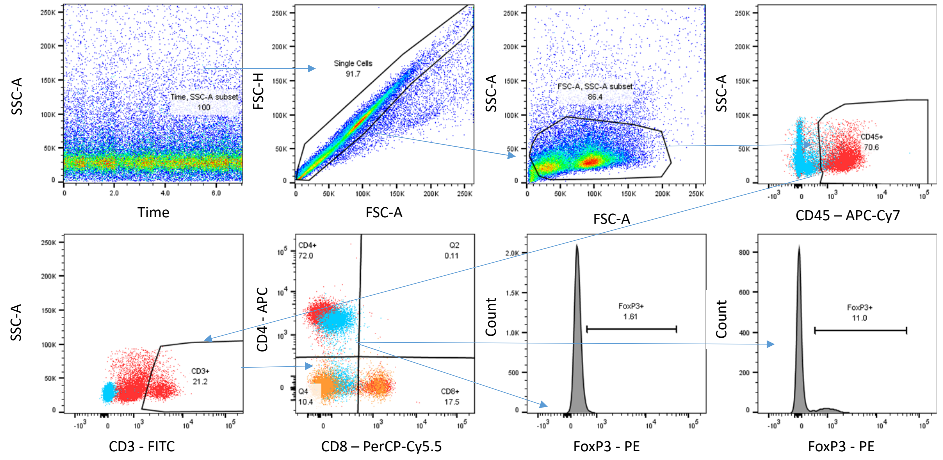


**B**

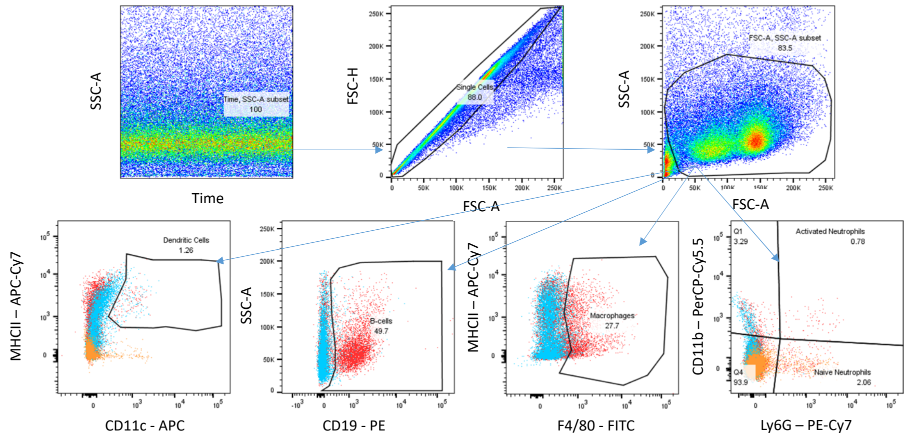
